# Molecular evolution of baculovirus IE1 protein, an essential multi-functional regulator of transcription and replication

**DOI:** 10.64898/2026.05.26.727872

**Authors:** Wujie Su, Chaoli Fang, Jiaru He, Wenbing Wang, Fanchi Li, Bing Li

## Abstract

Immediate-early protein IE1 is an essential protein of baculoviruses that is involved in transcription and replication. In the present study, we provide several lines of evidence for how ancient IE1 evolved into its current version. Using progressive truncations and site-directed mutations coupled with fluorescence microscopy, surprisingly, we showed that the previously identified nuclear localization sequence (NLS), basic domain II, was essential for sequence non-specific DNA binding, but not required for IE1 nuclear import, and demonstrated that, BmNPV IE1 (BmIE1) possesses not only a unique bipartite NLS that contains a monopartite NLS but also a cryptic non-canonical NLS that is correlated with sequence non-specific DNA binding. The non-canonical NLS alone was sufficient to launch infection. We also found that N-terminally truncated IE1 can enter the nucleus through its sequence non-specific binding ability. Remarkably, the monopartite NLS, when fused to the N-terminus of EGFP, can form a novel NLS that is also functional in a mammalian cell line. Moreover, residues 58 to 151 of BmIE1 were shown to be dispensable, and residue 152 was found to be critical for launching a productive infection. To gain insight into how ancient IE1 acquired its multi-functionality, we reorganized the N-terminal 23 aa and 132 aa of BmIE1 with EGFP in a variety of manners and found that both fragments are separable and transferable. Notably, we observed that the nuclear levels of BmIE1 should reach certain thresholds to initiate infection. Additionally, we found that BmNPV could launch infection more efficiently in a BmN cell line over another BmN cell line by increasing transcription levels of immediate early genes. Collectively, these findings suggest a hypothesis where ancient IE1 might have evolved the two additional NLSs and acquired the N-terminal 132 aa through gene fusion so as to reach infection-initiating thresholds at a faster pace.

**Author Summary:** Basic domain II was previously shown to be the NLS of AcMNPV IE1 protein. In the present study, we demonstrated that it is essential for sequence non-specific DNA binding, but not required for IE1 nuclear import, and found that BmNPV IE1 (BmIE1) possesses not only a unique bipartite NLS that contains a monopartite NLS but also a cryptic non-canonical NLS that is correlated with sequence non-specific DNA binding. We also found that N-terminally truncated BmIE1 can enter the nucleus through its sequence non-specific binding ability. Remarkably, the monopartite NLS, when fused to the N-terminus of EGFP, can form a novel NLS that is also functional in a mammalian cell line. Moreover, the domains of BmIE1 was shown to be separable and transferable. We also demonstrated that higher expression of functionally impaired BmIE1 achieved by higher MOIs or higher transfection efficiency can partially complement its compromised functions. Consistently, we found that BmNPV could launch infection more efficiently in a BmN cell line over another BmN cell line by increasing transcription levels of immediate early genes.

## 1 Introduction

Baculoviruses are pathogenic to invertebrates and characterized by a large circular double-stranded DNA genome that encodes 100-200 genes [1, 2]. Based on their phase of expression, these genes can be divided mainly into immediate-early, delayed-early, late and very late genes. Immediate early genes are critical for initiating viral replication in that their promoters can be recognized by host RNA polymerase II and thus can be expressed immediately after the viral genome enters host nucleus. By contrast, the expression of non-immediate-early genes is dependent on the products of immediate early genes. *Autographa californica* multicapsid nuclear polyhedrosis virus (AcMNPV) is the best-characterized baculovirus that has a wide host range, and *Bombyx mori* nuclear polyhydrosis virus (BmNPV), its most closely related relative, with its genome over 90% identical to about three-quarters of the AcMNPV genome[3], is infectious to the silkworm, *Bombyx mori*. Multiple homologous regions (hrs), putative origins of viral DNA replication, are found dispersed throughout both genomes [4, 5]. The hrs can also function as transcriptional enhancers [6]. The hrs are characterized by a series of ∼75-bp DNA repeats, with each containing a 30-bp imperfect palindrome with an *Eco*RI site at its core[4]. The immediate early protein IE1 binds as a dimer to the ∼28-bp im-perfect palindrome (28-mer) to exert its critical roles in transcription stimulation and DNA replication[7]. By its binding to hrs, baculovirus IE1 can trigger focal distribution in the nucleus of plasmid-transfected cells[8].

Both BmNPV IE1 (BmIE1) and AcMNPV (AcIE1) have been shown to be essential for launching a successful infection [9–12]. Consistent with its multi-functionality, AcIE1 has been shown to contain multiple functional domains (Fig.1 A). The N-terminal 23 residues of AcIE1 are required for origin-specific DNA replication and AcMNPV propagation, but not for DNA-binding-dependent transcriptional activation[13]. AcIE1 contains one transcriptional stimulatory domain from residues 8 to 118 and another one from 168 to 222[14, 15]. The two stimulatory domain is separated by a highly conserved basic Domain I (residues 152 to 161) that is required for hr enhancer DNA binding and hr-dependent transactivation[16]. In addition to a helix-loop-helix dimerization motif[15], a small basic domain from residues 534 to 538 (termed as basic domain II) that is required for DNA binding and nuclear import is found at its C terminus[17]. However, a recent study using truncated mutants of BmIE1 showed that IE1 (158–208) was a major nuclear localization element, and that IE (11–157) and IE1 (539–559) were minor nuclear localization elements [18]. Moreover, another study based on site-directed mutation of BmIE1 demonstrated that a domain spanning basic domain I served as a bipartite NLS of BmIE1[19]. This was inconsistent with a previous study demonstrating that the effect of deleting the entire basic domain I on nuclear localization was minimal[17], and thus further investigation is needed.

**Fig. 1.**
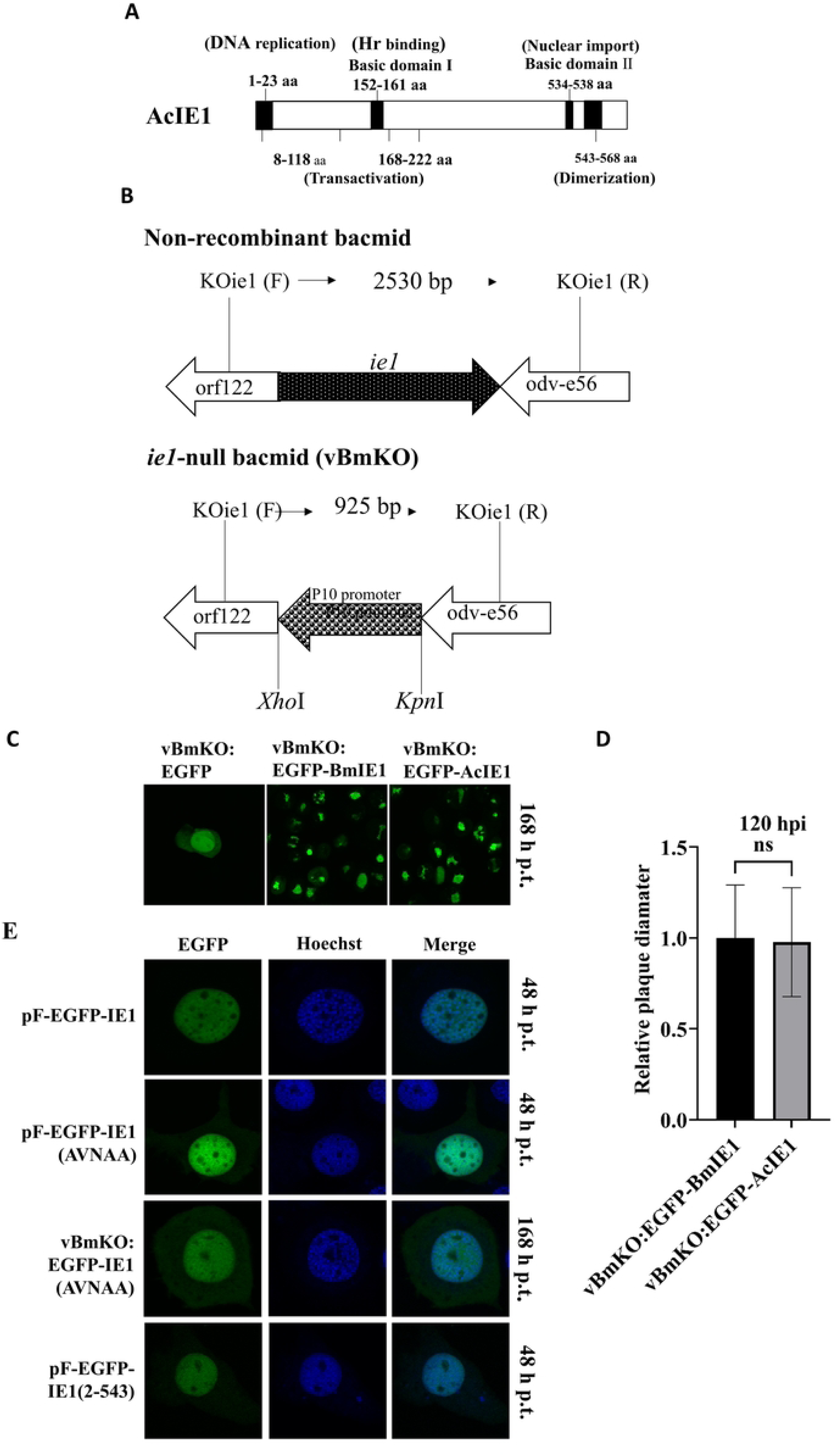
Generation of *ie1*-null BmNPV bacmid and Effects of basic domain II-mutated BmIE1 on subcellular localization. **A,** The previously identified functional domains of AcIE1. **B,** Genomic organizations of the *ie1* regions of wild-type and *ie1*-null BmNPV bacmids (vBmKO). **C,** Infectivity of repaired vBmKO. BmN cells were transfected with vBmKO: EGFP, vBmKO: EGFP-BmIE1 or vBmKO: EGFP-AcIE1, and were observed for signs of infection at 168 h p.t.. **D,** Relative plaque diameter. Twenty plaques were selected for each virus and their diameters were measured with Photoshop CS3. The ordinate shows the average diameter of vBmKO: EGFP-AcIE1 relative to that of vBmKO: EGFP-BmIE1. The bars represent standard deviations. **E,** Subcellular localization of basic domain II-mutated IE1 and dimerization domain-deleted IE1 in plasmid- and bacmid- transfected BmN cells. BmN cells were transfected with either plasmids or bacmid, fixed at 48 h p.t. (168 h p.t. for bacmid), stained with Hoechst 33342 and examined with a confocal microscope. Magnification, X400.

Small nuclear proteins can enter the nucleus through nuclear pore complexes via passive diffusion, whereas large nuclear proteins (over 40 Kda) should contain a NLS that can be recognized and bound by nuclear transport receptors for their active nuclear import [20]. Classical NLS motifs can be divided into two types: monopartite NLSs that consist of a single stretch of basic amino acids, and bipartite NLSs that possess two stretches of basic amino acids that are separated by a linker region[21]. The simian virus 40 (SV40) T-antigen NLS (126PKKKRKV132) is the canonical monopartite NLS[22]. In addition, there are also non-classical NLSs that are hard to be predicted for their lack of typical features. Some proteins can also enter the nucleus through protein-protein interactions. For example, AcMNPV P143, a cytoplasmic helicase, can enter the nucleus via its interaction with LEF-3[23].

In the present study, we found that BmIE1 can enter the nucleus via 3 NLSs: one bipartite NLS, one monopartite NLS and one non-canonical NLS. Interestingly, for some truncated IE1 mutants, they can localize to the nucleus even by taking advantage of its non-specific DNA binding activity. Moreover, we reorganized the N-terminal 23 aa and 132 aa of BmIE1 with EGFP in a variety of manners and found that both fragments are separable and transferable. For example, we inserted EGFP between residues 132 and 133 of BmIE1 or transferred the N-terminal 132 aa to the C-terminus of EGFP that is fused to N-132 aa-truncated IE1 at the C-terminus. Remarkably, both mutants were capable of launching infection. Notably, we found that higher levels of impaired IE1 achieved by higher MOIs or higher transfection efficiency could partially compensate for compromised function. Consistently, we also found that BmNPV could launch infection more efficiently in a BmN cell line over another BmN cell line by increasing transcription levels of immediate early genes, and this might serve as a force that drove the evolution of IE1. Taken together, these findings provide insight into how the multi-functional protein IE1 evolved.

## 2 Results

### 2.1 Basic domain II is not required for IE1 nuclear import

To be able to investigate the effects of IE1 mutations in the context of infection, we first generated an *ie1*- knockout mutant. To rule out the possibility that homologous recombination might occur between the retaining *ie1* sequences at the *ie1* locus and mutant *ie1* that is reinserted at the polyhedrin locus, the whole *ie1* open reading frame was deleted using a selection marker-free method previously described [12]. As the promoter of *ac146* (*orf126* in BmNPV), an essential late gene[24], is contained in the *ie1* sequence, the *p10* promoter is inserted (Fig. 1B). An *ie1*-null mutant named vBmKO was generated (Fig.1B). BmIE1, possessing two additional residues, has 95.55% amino acid identity with AcIE1 (S1Fig.). Bacmid transfection (Fig.1C) and plaque diameter analysis (Fig.1D) showed that AcIE1 can effectively complement BmIE1 despite their differences.

A previous study based on biochemical fractionation showed that a dimeric nuclear element, basic domain II (KVNRR), is required for IE1 nuclear import[17]. Unexpectedly, in our plasmid transfection assay, EGFP fused with basic domain II-mutated IE1 (EGFP-IE1 (AVNAA)) still localized to the nucleus, with a weak fluorescent signal observed in the cytoplasm (Fig. 1E). A similar result was observed with bacmid transfection of BmN cells (Fig. 1E). Moreover, in bacmid-transfected BmN cells, no signs of infection or no focal loci were observed (Fig.1E). Additionally, EGFP-IE1 (2-543), with a C-terminal truncation (residues 544 to 584) that disrupts IE dimerization, was still found to localize to the nucleus (Fig.1E). These findings suggest that basic domain II is not necessary for IE1 nuclear import and that IE1 contains at least one NLS that is independent of dimerization, but that basic domain II is essential for infection and is required for formation of focal loci.

### 2.2 Residues 135 to 163 constitute a bipartite NLS that contains one monopartite NLS

To map IE1 NLSs, a progressive truncation method was used, and a series of IE1 truncations fused with EGFP were constructed. EGFP-IE1 fusions, with IE1 residues from 2 to 134 deleted, were all found predominantly in the nucleus of plasmid-transfected BmN cells (S2 Fig.). EGFP-IE (135-584) was found mainly in the nucleus, whereas EGFP-IE (136-584) localized predominantly to the cytoplasm (Fig. 2A). Similarly, EGFP-IE (K135A) and EGFP-IE (R136A) localized predominantly to the cytoplasm, whereas EGFP-IE (K137A) were distributed both within the nucleus and cytoplasm (Fig. 2A). These results indicated that the KRK motif played a critical role in IE1 nuclear localization. Moreover, we found that mutation of Lys135 to Arg (RRK) or mutation of Lys135 and Arg136 to His (HHK) could partially restore IE1 nuclear import, whereas mutation of Arg136 to Lys (KKK) failed to effectively restore IE1 nuclear targeting (Fig. 2A). To determine the whole NLS, N-terminal IE1 residues varying in the C-terminus were fused with EGFP. Relatively strong fluorescent signal was found in the cytoplasm for EGFP-IE1 (2-144) and EGFP-IE1 (2-155) (Fig. 2B), whereas EGFP-IE1 (2-163) and EGFP-IE1 (2-193) were localized predominantly to the nucleus, with no marked difference in their nuclear import (Fig. 2B). Moreover, EGFP-IE1 (2-163) was fused with *Bombyx mori* actin A3, a cytoplasmic protein, and the resulting fusion protein EGFP-IE1 (2-163)-A3 was found completely in the nucleus (Fig. 2B). These results suggested that IE1 contained an NLS in its N-terminal 163 residues. As the N-terminally truncated IE1 mutants showed that the N-terminal 134 residues were dispensable for IE1 nuclear import, we speculate that residues from 135 to 163 forms an NLS of IE1. To test this, IE1 (136-163) and IE1 (135-163) were fused with EGFP. As expected, IE1 (136-163) lacking Lys135 failed to direct EGFP completely to the nucleus (Fig.2 B). By contrast, IE1 (135-163) containing the entire intact NLS domain was able to direct EGFP and A3 completely to the nucleus (Fig. 2B). Moreover, IE1 (135-163) was also found to direct P143 completely to the nucleus (S3 Fig.). Given that residues 154 to 163 (KNKLKPKYKK) are rich in basic residues, we hypothesized that, combined with the motif KRK, the amino acid sequence constitutes a bipartite NLS of BmIE1. To test this hypothesis, we constructed a series of mutations in residues from 138 to 163 using point mutations. Unexpectedly, none of these mutations had a considerable effect, although weak fluorescent signal was observed in the cytoplasm for 5 mutations (LDE to AAA, YLD to AAA, K158A/P159A, K162A and K163A) (S4 Fig.).

**Fig. 2.**
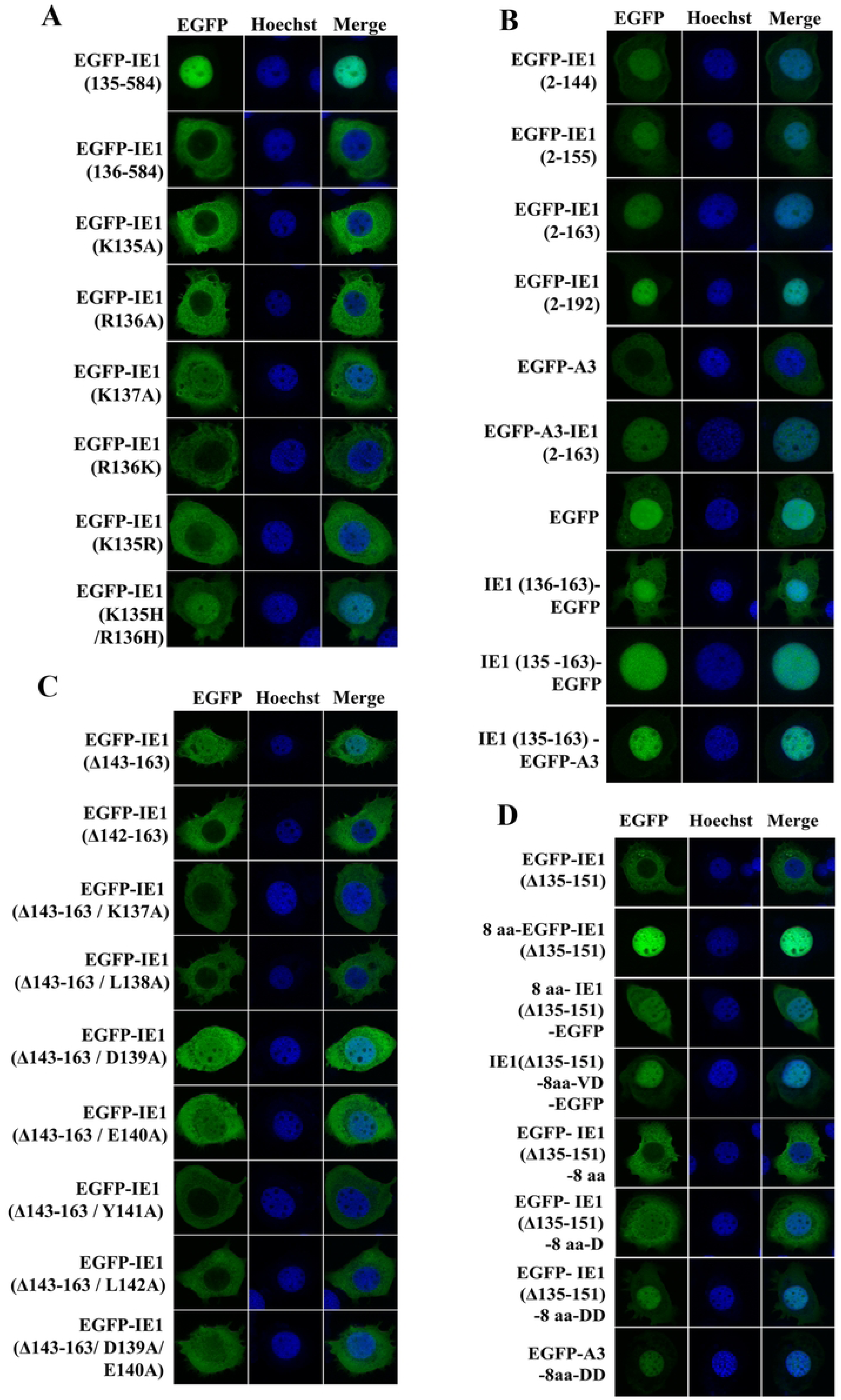
Residues 135 to 163 of BmIE1 constitute a bipartite NLS that contains one monopartite NLS in BmIE1. BmN cells were transfected with plasmids, fixed at 48 h p.t., stained with Hoechst 33342 and examined with a confocal microscope Magnification, X400. **A,** Identification of the critical role of K135, R136 and K137 in IE1 nuclear import. **B,** Identification of a bipartite NLS (residues 135 to 163). **C, D,** Identification of a monopartite NLS (residues 135 to 143).

Deletion of the entire basic domain I was previously shown to have a minimal effect on IE1 localization[17]. However, in our study, deletion of the entire basic domain I (residues 152 to 163) appeared to impair but not abolish nuclear targeting (S5 Fig.), indicating that basic domain I indeed plays a role in IE1 nuclear import but is not fully responsible. To identify remaining residues that are involved in IE1 nuclear targeting, we continued to delete residues N-terminal to basic domain I. No considerable effect was observed until residues 142 to 163 was deleted (Fig. 2C and S5 Fig.). If residues 135 to 163 form a bipartite NLS, which requires two stretches of basic residues, deletion of the entire basic domain I and residues N-terminal to it should abrogate its ability to direct IE1 nuclear targeting. These findings suggest that Leu142 might be an essential residue of another NLS.

To test this possibility, we first determined the importance of residues from 137 to 142 using point mutations. Remarkably, EGFP-IE1 (Δ143-163/L142A), EGFP-IE1 (Δ143-163/Y141A), EGFP-IE1 (Δ143-163/L138A), EGFP-IE1 (Δ143-163 /K137A), localized predominantly to the cytoplasm (Fig. 2C), whereas EGFP-IE1 (Δ143-163/ E140A), EGFP-IE1 (Δ143-163/ D139A) and EGFP-IE1 (Δ143-163/ D139A/ E140A) were distributed both in the nucleus and cytoplasm (Fig. 2C). Given that EGFP-IE (K135A) and EGFP-IE (R136A) localized predominantly to the cytoplasm (Fig. 2A), these findings suggest that the 8 residues, KRKLDEYL (referred to as 8 aa hereafter), might constitute a monopartite NLS. To confirm whether the 8 aa domain can direct a protein to the nucleus, we fused the 8 aa to the EGFP-IE1(Δ135-151) fusion protein at multiple sites. As expected, 8 aa-EGFP-IE (Δ135-151) localized completely to the nucleus, and 8 aa-IE1 (Δ135-151)-EGFP and IE1 (Δ135-151)-8 aa-VD-EGFP localized predominantly to the nucleus (Fig. 2D). Unexpectedly, EGFP-IE (Δ135-151)-8aa localized predominantly to the cytoplasm (Fig. 2D). Notably, EGFP-IE1 (Δ135-151)-8 aa-D and EGFP- IE1 (Δ135-151)-8 aa-DD in which one and two additional Asp residues are respectively attached to the C-terminus of the 8 aa domain regained the nuclear targeting ability (Fig. 2D). Moreover, EGFP-A3-8 aa-DD (Fig.2D) and EGFP-P143-8 aa-DD (S3 Fig.) also localized predominantly to the nucleus. These results indicate that the 8 aa domain is a genuine monopartite NLS and that Asp might be not essential but can augment nuclear targeting.

As 8 aa-EGFP-IE (Δ135-151) localized completely to the nucleus, we also investigated which residues of EGFP can improve nuclear targeting of EGFP-IE (Δ135-151)-8 aa by attaching N-terminal residues of EGFP to the C-terminus of the 8 aa domain. Unexpectedly, unlike fusion proteins with Asp-attached 8 aa-monopartite NLS, EGFP-IE1 (Δ135-151)-8 aa-EGFP (2-4), or EGFP-IE1 (Δ135-151)-8 aa-EGFP (2-9) or even EGFP-IE1 (Δ135-151)-8 aa-EGFP (2-239) were all found to localize predominantly to the cytoplasm (Fig. 3A). Given that IE1(Δ135-151)-8 aa-VD-EGFP localized predominantly to the nucleus (Fig. 2D) and it is different from EGFP-IE1 (Δ135-151)-8 aa-EGFP (2-239) in that its 8 aa domain is connected with EGFP (2-239) by two additional residues Val and Asp (translated from the SalI restriction site GTCGAC), we hypothesized that the 8 aa domain can have a strong activity either when it has one or two Asp residues attached to its C-terminus or when it has a specific sequence context (i.e., location-specific or conformation-specific). To test this possibility, we replaced the N-terminal EGFP of EGFP-IE1 (Δ135-151)-8 aa-EGFP (2-239) with 8 aa-EGFP (2-5), surprisingly, the resulting fusion protein 8 aa-EGFP (2-5)-IE1 (Δ135-151)-8 aa-EGFP (2-239) gained partial nuclear import (Fig. 3A), indicating that the 8 aa domain with N-terminal residues 2 to 5 of EGFP attached indeed has a nuclear targeting ability when it is located at the N-terminus of the fusion protein. Moreover, when we attached more N-terminal residues of EGFP to the 8 aa domain, the nuclear targeting ability of the 8 aa domain was increasingly augmented (S6 Fig.). The nuclear targeting ability of the 8 aa domain nearly reached a level achieved by attaching the full-length EGFP (2-239) when we attached 11 N-terminal residues (i.e., 2-12) of EGFP (Fig. 3A), indicating that the 11 N-terminal residues of EGFP are the main residues responsible for improving the nuclear targeting ability of the 8 aa domain. To rule out the possibility that the 8 aa domain at the C terminus of the fusion protein might have an effect, we deleted it and found that 8 aa-EGFP (2-12)-IE1 (Δ135-151)-EGFP (2-239) has the same nuclear targeting ability as 8 aa-EGFP (2-12)-IE1 (Δ135-151)-8aa-EGFP (2-239) (Fig. 3A). Moreover, 8 aa-EGFP-A3 was evenly distributed between cytoplasm and nucleus (Fig.3A), whereas 8 aa-EGFP-P143 localized predominantly to the nucleus (S3 Fig.). Taken together, these findings suggest that the nuclear targeting ability of the 8 aa domain can be improved by the N-terminal residues of EGFP and that this effect is location-specific.

**Fig. 3.**
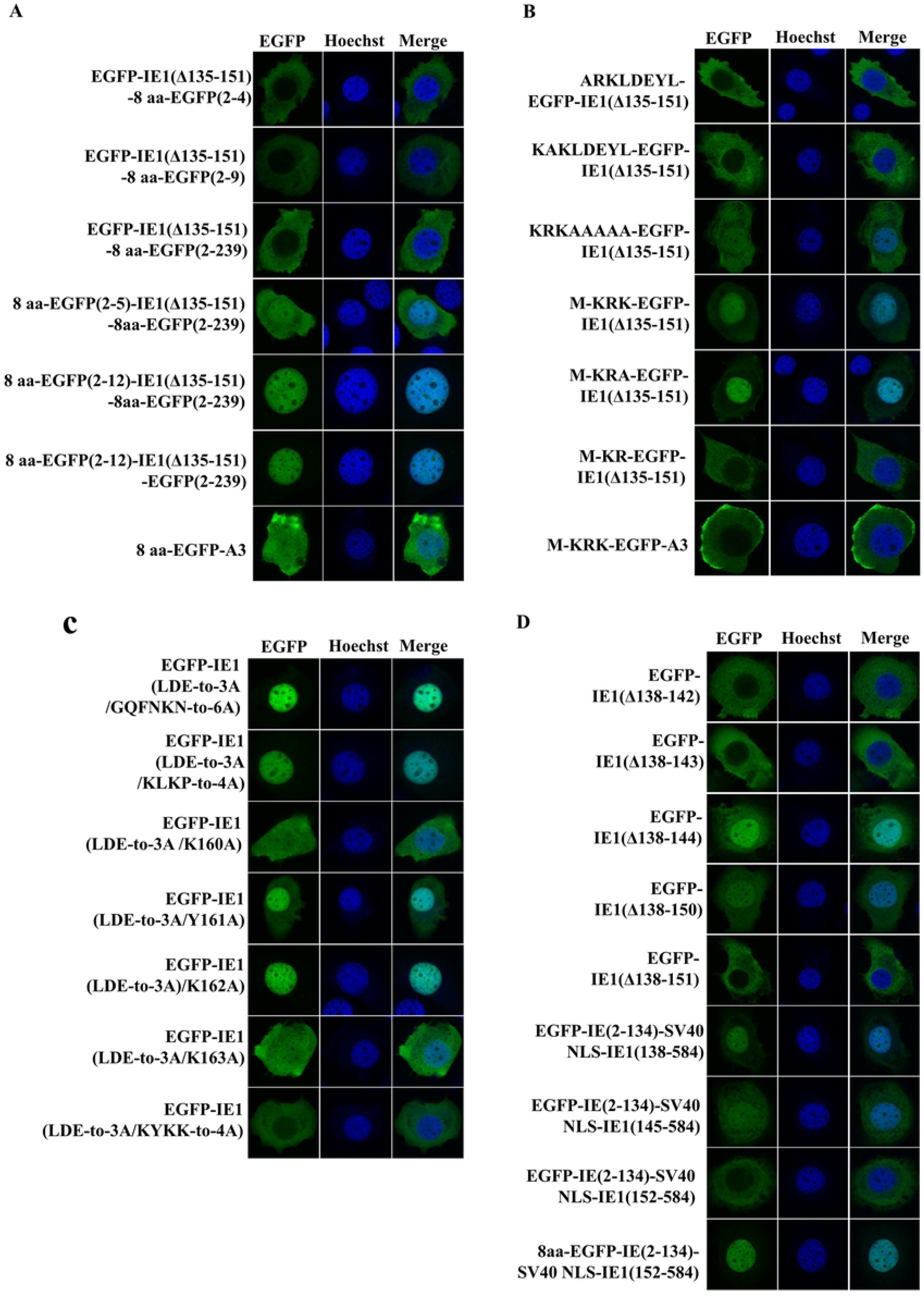
Identification of a novel NLS formed by the 8 aa domain and N-terminal residues of EGFP as well as critical basic residues in the bipartite NLS. BmN cells were transfected with plasmids, fixed at 48 h p.t, stained with Hoechst 33342 and examined with a confocal microscope Magnification, X400. **A,** Identification of critical N-terminal residues of EGFP involved in the 8 aa-EGFP NLS. **B,** Unique features of the newly identified NLS: M-KRK-EGFP. **C,** Identification of critical residues in the bipartite NLS. D, The long linker length of the bipartite is critical for the nuclear targeting ability of the 8 aa monopartite NLS.

Why the N-terminal residues of EGFP can augment the nuclear targeting of the 8 aa domain is intriguing. It is possible that the basic residue Lys4 and the two consecutive acidic residues Glu6 and Glu7 of EGFP play an important role. Alternatively, the N-terminal residues of EGFP might form a NLS with the KRK motif in the 8 aa domain. To test this possibility, we constructed a series of point mutations. As expected, mutation of KRK to ARK (ARKLDEYL-EGFP-IE1 (Δ135-151)) or KAK (KAKLDEYL-EGFP-IE1 (Δ135-151)) in the 8 aa domain caused loss of nuclear targeting abilities of the fusion proteins (Fig. 3B). However, none of the other residues in the 8 aa domain were essential for nuclear targeting (Fig. 3B and S7 Fig.). In particular, the mutant KRKAAAAA-EGFP-IE1 (Δ135-151) still retained its nuclear targeting ability (Fig.3B). As a monopartite NLS, only Asp and Glu in the 8 aa domain are not essential for nuclear targeting (Fig. 2A and 2C). These results indicate that the KRK motif might form a NLS with N-terminal residues of EGFP. Since the last 5 residues in the 8 aa domain were found to be not essential, we deleted them to determine whether the putative NLS still works. Surprisingly, the resulting fusion protein M-KRK-EGFP-IE1 (Δ135-151) still retained its nuclear targeting ability (Fig. 3B). Moreover, although M-KRA-EGFP-IE1 (Δ135-151) also entered the nucleus (Fig. 3B), M-KR-EGFP-IE1 (Δ135-151) appeared to lose its nuclear targeting ability (Fig. 3B), indicating that the linker length is also critical (a feature typical of bipartite NLSs). We also substituted KR for other basic residues and found combinations of other residues did not have the same effect as the KR combination (S7 Fig.). Unexpectedly, M-KRK-EGFP-A3 and M-KRK-EGFP-P143 was found to localize predominantly to the cytoplasm (Fig. 3B and S3 Fig.). Taken together, these results suggest that the KRK motif in the 8 aa domain form a NLS with the N-terminal residues of EGFP, and that the M-KRK motif forms a NLS with the fusion protein EGFP-IE1 (Δ135-151), although this nucleus targeting ability is specific to IE1 among the 3 proteins we tested.

Since the 8 aa domain contained in the putative 29 aa bipartite NLS can greatly mask the effect of point mutations (S4 Fig.), we deactivated the 8 aa monopartite NLS by mutating LDE to AAA to determine which residues are critical for the putative bipartite NLS. The 6 residues GQFNKN and the four residues KLKP were found to contribute little if any to nuclear targeting ability (Fig. 3C). Mutation of Lys162 to alanine also had little effect (Fig. 3C). Tyr161 was found to play some role (Fig. 3C). Lys160 and Lys163 were shown to play a major role in the nuclear targeting ability of the 29 aa bipartite NLS and mutation of the two residues and the two residues between them to alanine rendered the resulting fusion protein localize predominantly to the cytoplasm (Fig. 3C). Taken together, these results show that the putative 29 aa NLS is a genuine bipartite NLS with a 22 aa linker that separates the KRK motif and the KYKK motif.

Bipartite NLSs normally have linkers of 9 to 12 residues, whereas the bipartite NLS identified here possesses a linker of 22 aa. To determine whether a linker as long as 22 aa is necessary for the bipartite NLS, we conducted a progressive deletion of the linker. Surprisingly, when we deleted the first 5 or 6 residues of the linker (EGFP-IE1 (Δ138-142) and EGFP-IE1 (Δ138-143)), the nuclear targeting ability of the bipartite NLS appeared to be abolished (Fig. 3D). However, deletion of more residues (7 to 13) from the linker rendered the NLS regain nuclear targeting ability (Fig. 3D and S8 Fig.). Interestingly, deletion of the first 14 or more residues, which made the linker 8 aa or shorter in length, abrogated the nuclear targeting ability of the NLS (Fig.3D and S8 Fig.), indicating that the bipartite NLS itself requires a linker of 9 aa or more. Since a linker of 9 aa is sufficient, it is interesting to investigate why BmIE1 evolved a linker of 22 aa. To answer this question, we replaced residues 135 to 137 (i.e., the motif KRK), residues 135 to 144 and residues 135 to 151 with the classic monopartite NLS: SV40 NLS. Remarkably, the nuclear targeting abilities of these fusion proteins decreased as the SV40 NLS approached basic domain I (Fig. 3D). Moreover, the abolished nuclear targeting ability of EGFP-IE (2-134)-SV40 NLS-IE1 (152-584) could be repaired by attaching the 8 aa domain to the N-terminus of EGFP. Taken together, these results indicate that the 29 aa bipartite NLS might have evolved the long linker to protect the nuclear targeting ability of the 8 aa monopartite NLS.

### 2.3 IE1 contains a non-canonical NLS that is correlated with sequence non-specific DNA binding

Although vBmKO: EGFP-IE1 (Δ135-137) failed to spread in transfected BmN cells, with EGFP-IE1 (Δ135-137) localized predominantly to the cytoplasm (Fig. 4A), focal loci were seen both in the cytoplasm and nucleus of BmN cells transfected with an hr3-containing plasmid pF-hr3-ie1p-EGFP-IE1 (Δ135-137) (Fig.4A). These findings suggest that BmIE1 might still have a cryptic NLS because deletion of the KRK motif deactivates both the 29 aa bipartite NLS and the 8 aa monopartite NLS. To test this possibility, a series of N-terminal truncations were constructed. Interestingly, while EGFP-IE1 (202-584) and EGFP-IE1 (298-584) localized predominantly to the cytoplasm, EGFP-IE1 (373-584) and EGFP-IE1 (387-584) were distributed mainly in the nucleus (Fig. 4B). However, deletion of more residues, i.e., EGFP-IE1 (396-584) and EGFP-IE1 (426-584), appeared to result in an impaired nuclear targeting ability (Fig. 4B). It is possible that residues from 373 to 395 might form an essential part of an NLS. However, the fusion protein IE1 (373-437)-A3-EGFP was found mainly in the cytoplasm (Fig. 4C). Moreover, even IE1 (373-584)-A3-EGFP failed to localize predominantly to the nucleus (Fig. 4C). Taken together, these findings suggest that the C-terminus of BmIE1 (residues 373 to 584) contains a unique NLS that can function efficiently only when the size of the fusion protein is proper.

**Fig. 4.**
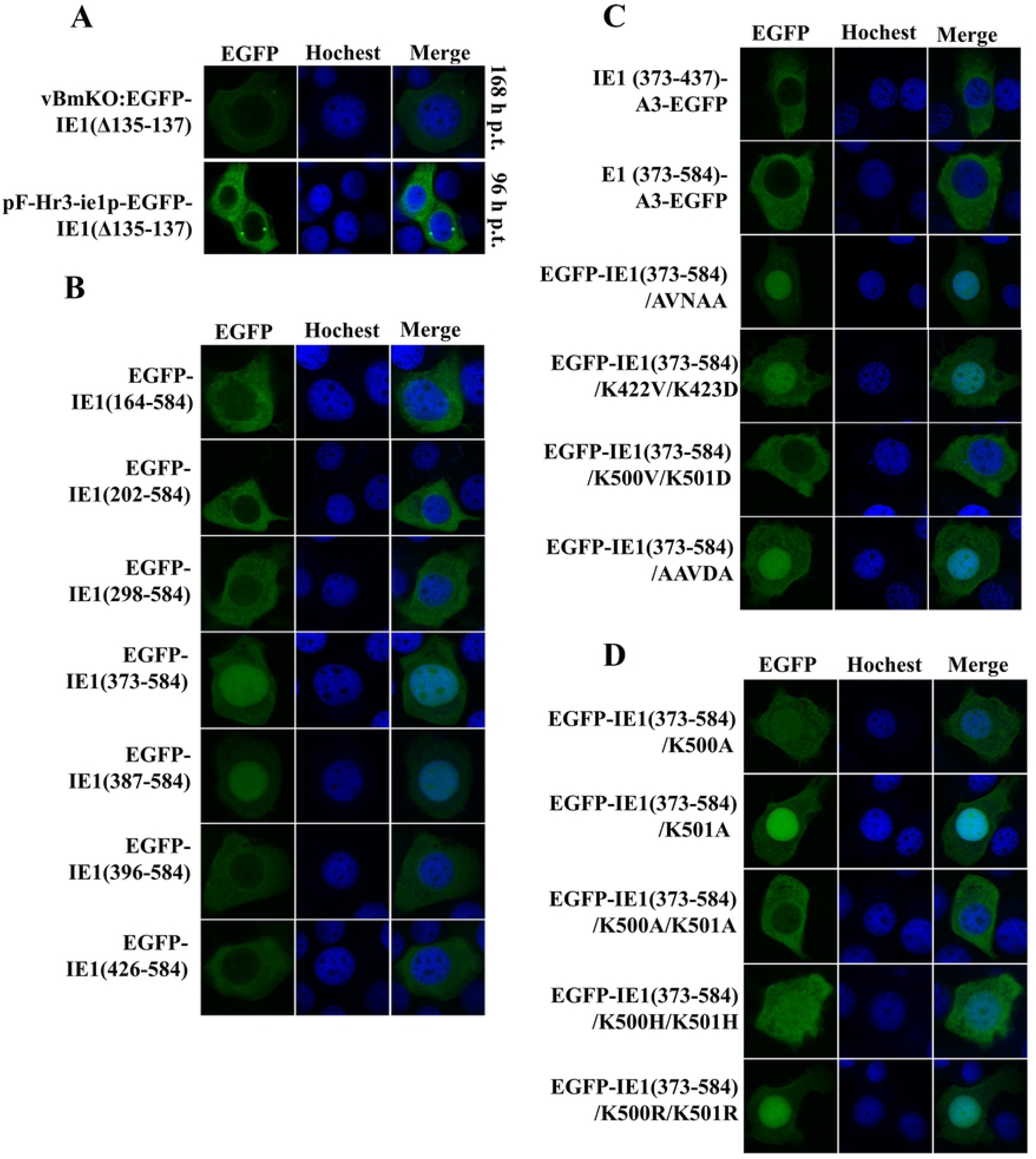
Identification of a non-canonical NLS in BmIE1. BmN cells were transfected with plasmids, fixed at 48 h p.t., stained with Hoechst 33342 and examined with a confocal microscope Magnification, X400. **A,** Formation of focal loci by EGFP-IE1 (Δ135-137) in the nucleus. Although vBmKO: EGFP-IE1(Δ135-137) failed to launch infection at 168 h p.t., in the nucleus of BmN cells transfected with pF-Hre-ie1p-EGFP-IE1 (Δ135-137), focal loci could be observed at 96 h p.t.. **B,** Identification of truncated IE1 that can still enter the nucleus. **C,** Identification of a non-canonical NLS (K500 and K501). **D,** Effects of replacements of K500 and K501 with other residues on subcellular localization.

Since EGFP-IE1 (373-584) localized predominantly to the nucleus, we used this fusion protein as a backbone to identify residues that are critical to the cryptic NLS. Inspection of the amino acid sequence of the C-terminus of BmIE1 revealed 3 regions with at least 2 consecutive Lysine, i.e., residues 422 to 423 (KK), residues 457 to 461 (KKVKK), and residues 500 to 501 (KK). To determine whether these lysine are important for nuclear targeting, we conducted site-directed mutation on these residues of EGFP-IE1 (373-584). Moreover, we also investigated whether mutated basic domain II has an effect on the nuclear targeting ability of EGFP-IE1 (373-584). As can be seen from Fig. 4C, EGFP-IE1 (373-584)-K422V/K423D, EGFP-IE1 (373-584)-KKVKK-to-AAVDA, EGFP-IE1 (373-584)-KVNRR-to-AVNAA localized predominantly to the nucleus, whereas EGFP-IE1 (373-584)-K500V/K501D was distributed predominantly in the cytoplasm, indicating that Lys500 and Lys501 are critical for nuclear targeting.

To further determine whether both lysine are critical and whether they can be substituted for other basic residues, a series of point mutations were constructed. K501A mutation appeared to have little if any effect on nuclear targeting (Fig. 4D), whereas K500A mutation considerably impaired nuclear targeting (Fig. 4D). Double mutation (K500A/K5001A) resulted in more impaired nuclear targeting than the K500A single mutation (Fig. 4D), indicating that, although Lys500 is sufficient for nuclear import, Lys501 can partially complement Lys500. Moreover, basic residues work better than alanine in this motif, and EGFP-IE1 (1-372)-K500R/K501R possesses a better nuclear targeting ability than EGFP-IE1 (373-584)-K500H/K501H (Fig. 4D).

To investigate whether these basic residues are critical in the context of baculovirus infection, we re-introduced *egfp-ie1* fusion genes harboring mutations on these residues to the *polyhedrin* locus of vBmKO under the control of the *ie1* promoter. As with vBmKO: EGFP-IE1 (KVNRR-to-AVNAA), in BmN cells transfected with bacmids including vBmKO: EGFP-IE1 (K422V/K423D), vBmKO: EGFP-IE1 (KKVKK-to-AAVDA) and vBmKO: EGFP-IE1 (K500V/K501D), no signs of infection or no focal loci were observed (Fig. 5A), indicating that these basic residues are essential and that they are directly or indirectly involved in hr binding. In support of this, in mitotic cells, EGFP-IE1 (Δ135-137)/ (K422V/K423D), EGFP-IE1 (Δ135-137)/ (KKVKK-to-AAVDA) and EGFP-IE1 (Δ135-137)/ (K500V/K501D) failed to bind to condensed chromosomes, whereas binding of EGFP-IE1 (Δ135-137) to condensed chromosomes was observed (S9 Fig.).

**Fig. 5.**
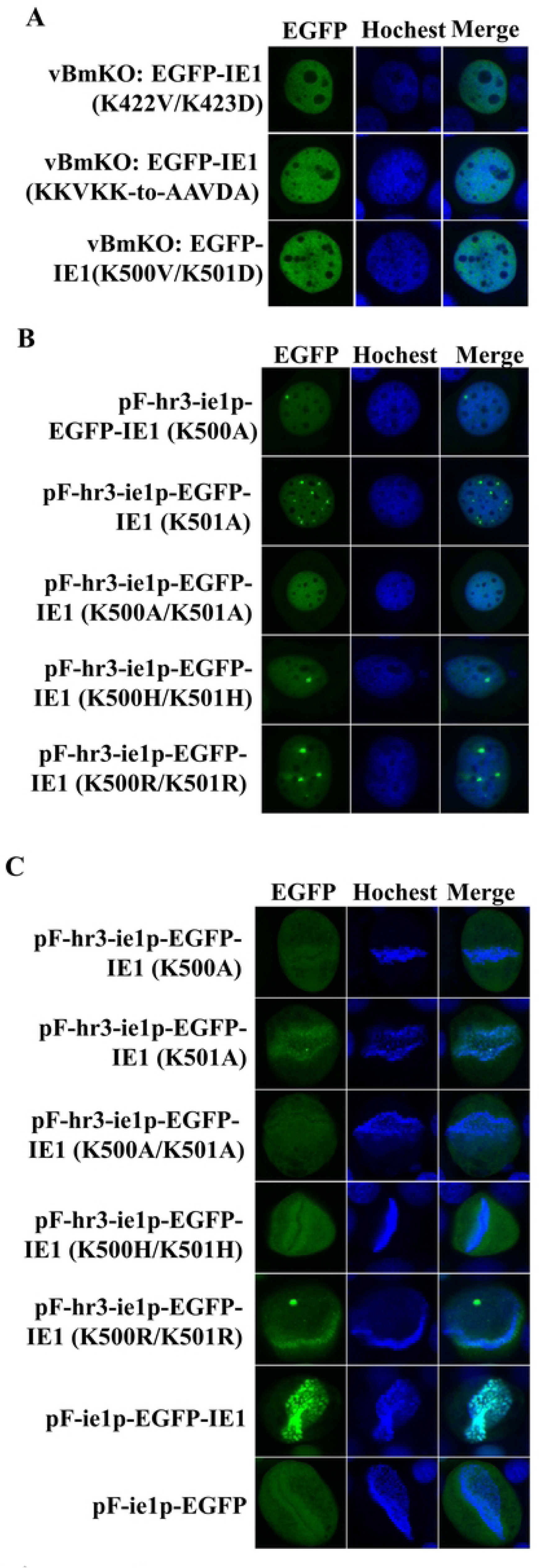
The non-canonical NLS is correlated with sequence non-specific DNA binding. BmN cells were transfected with either plasmids or bacmid, fixed at 48 h p.t. (168 h p.t. for bacmid), stained with Hoechst 33342 and examined with a confocal microscope Magnification, X400. **A**, infectivity of vBmKO repaired by EGFP-IE1 (K422V/K423D), EGFP-IE1 (KKVKK-to-AAVDA) and EGFP-IE1 (K500V/K501D). No signs of infection or focal loci were observed for all these repaired viruses at 168 h p.t.. **B,** Involvement of the non-canonical NLS in hr binding. The K500A/K501A double mutation results in failure to form focal loci. **C,** Involvement of the non-canonical NLS in sequence non-specific DNA binding. Mutations in the non-canonical NLS are correlated with an impaired ability to bind to condensed chromosomes in mitotic cells.

As Lys500 and Lys501 are also involved in nuclear targeting, we selected this motif for further analysis. To confirm whether the motif is related to hr binding, we constructed plasmids that contains an hr3 sequence upstream of the *ie1* promoter and the fusion genes were placed downstream. For all mutations, with the exception of the K500A/K5001A double mutation, focal loci were observed in plasmid-transfected BmN cells at 96 h p.t. (Fig. 5B). Moreover, in mitotic cells, binding of EGFP-IE1, EGFP-IE1 (K501A) and EGFP-IE1 (K500R/K501R) to condensed chromosomes was observed, whereas EGFP-IE1 (K500A), EGFP-IE1 (K500A/K501A) and EGFP-IE1 (K500H/K501H) appeared to have a similar localization with EGFP (Fig. 5C). Taken together, these findings suggest that Lys500 and Lys501 are indeed involved in hr binding and this ability appears to positively correlate with its capability to bind to condensed chromosomes (i.e., sequence non-specific DNA binding ability).

### 2.4 N-terminally truncated IE1 can enter the nucleus through its sequence non-specific DNA binding ability

Interestingly, at 96 h p.t., in BmN cells transfected with pF-ie1p-EGFP-IE1 (136-584), EGFP-IE1 (136-584) was observed to localize predominantly to the nucleus or the cytoplasm, or co-localize with condensed chromosomes in different cells (Fig. 6A). This special localization pattern was not specific to EGFP-IE1 (136-584). We found that EGFP-IE1 (164-584), a truncation mutant in which both the bipartite NLS and the monopartite NLS and the entire basic domain I (hr binding domain) were deleted, also had these 3 forms of localization in different cells (Fig.6B). However, when an additional mutation (KVNRR to AVNAA) was introduced to EGFP-IE1 (163-584), the resulting fusion protein EGFP-IE1 (164-584)/(KVNRR-to-AVNAA) no longer co-localized with condensed chromosomes and was not observed to localize predominantly to the nucleus (Fig. 6B). Moreover, as those cells with EGFP-IE1 (136-584) or EGFP-IE1 (164-584) localized predominantly to the nucleus were found always in pairs, we hypothesized that cytoplasmic EGFP-IE1 (136-584) and EGFP-IE1 (164-584) enter the nucleus by binding to condensed chromosomes during mitosis when the nuclear envelope break downs via their sequence non-specific DNA binding ability. As the KVNRR to AVNAA mutation abolished this ability, we speculated that, like the non-canonical NLS, basic domain II has a role in sequence non-specific DNA binding. To test this hypothesis, we co-transfected pF-ie1p-EGFP-IE1 (Δ135-137) with pF-ie1p-39K-DsRed into BmN cells. 39K is a nuclear baculovirus-encoded protein that has been shown to have a sequence non-specific DNA binding ability[25]. As expected, in non-dividing cells, EGFP-IE1 (Δ135-137) localized predominantly to the cytoplasm, whereas 39K-Dsred was found mainly in the nucleus (Fig. 6C). In dividing cells, both fusion proteins bound to condensed chromosomes (Fig. 6C). However, when the KVNRR to AVDAA mutation was introduced to EGFP-IE1(Δ135-137), the resulting fusion protein no longer co-localized with 39K-Dsred or condensed chromosomes in mitotic cells (Fig. 6C). Moreover, we found that, although both EGFP-IE1 (164-584) and EGFP-IE1 (164-584)-AVNAA were able to form focal loci in the nucleus by heterodimerizing with wild-type IE1 in infected BmN cells, after fixation by 4% paraformaldehyde, focal loci became diffuse in the nuclei of a great portion of BmN cells for the EGFP-IE1 (164-584)/AVNAA group(Fig.6D), indicating that both copies of intact KVNRR domain are required for the dimer to have a strong DNA binding ability. Taken together, these results confirm that it is sequence non-specific DNA binding ability that is responsible for helping N-terminally truncated IE1 enter the nucleus.

**Fig. 6.**
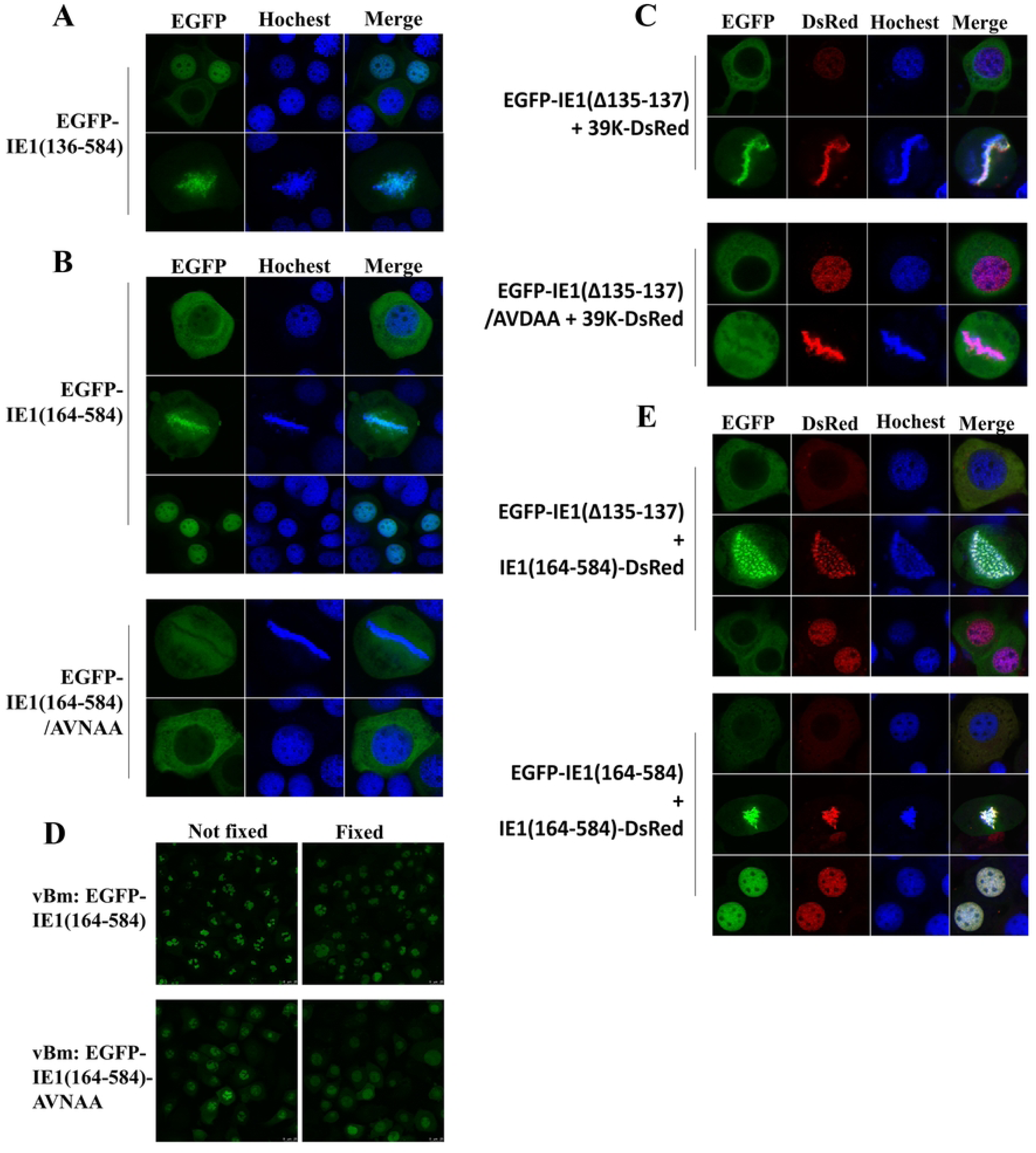
Truncated IE1 can enter the nucleus through its sequence non-specific DNA binding ability. BmN cells were transfected with plasmids, fixed at 96 h p.t., stained with Hoechst 33342 and examined with a confocal microscope Magnification, X400. **A,** Three forms of EGFP-IE1 (136-584) subcellular localization. **B,** Three forms of EGFP-IE1 (164-584) subcellular localization and two forms of EGFP-IE1 (164-584)/AVNAA subcellular localization. A mutation in basic domain II rendered the truncated IE1 lose its ability to bind to condensed chromosomes and the nuclear subcellular localization form was not observed. **C,** Co-localization of EGFP-IE1 (Δ135-137) with 39K-DsRed through its sequence non-specific DNA binding ability in mitotic cells. A mutation in basic domain II (KVNRR-to-AVDAA) rendered the truncated IE1 lose its ability to bind to condensed chromosomes and failed to co-localize with 39K-DsRed in mitotic cells. **D,** Truncated IE1 is more stable after mitosis. In BmN cells co-transfected with EGFP-IE1 (Δ135-137) and IE1 (164-584)-DsRed, red fluorescent signal was observed in the nucleus after mitosis, whereas the green fluorescent signal was observed in the cytoplasm, despite the observation that both protein bind to condensed chromosomes in mitotic cells. By contrast, EGFP-IE1 (164-584) co-localized with IE1 (164-584)-DsRed in all cases.

Although EGFP-IE1 (Δ135-137) was able to bind to condensed chromosomes (Fig. 6C), we rarely observed cells with EGFP-IE1 (Δ135-137) localized predominantly to the nucleus. To explore the mechanism by which EGFP-IE1 (Δ135-137) failed to remain in the nucleus after mitosis, we co-transfected pF-ie1p-EGFP-IE (Δ135-137) with pF-ie1p-IE1 (Δ1-163)-DsRed into BmN cells. As can be seen from Fig. 6E, in non-dividing cells, EGFP-IE (Δ135-137) and IE1 (164-584)-DsRed both localized to the cytoplasm, and in mitotic cells, they both bound to condensed chromosomes. However, in cells that had completed mitosis, EGFP-IE (Δ135-137) localized predominantly to the cytoplasm, whereas IE1 164-584)-DsRed was found primarily in the nucleus (Fig.6E). To determine whether it was the N-terminus of BmIE1 that was responsible for the difference, we then co-transfected pF-ie1p-EGFP-IE (164-584) with pF-ie1p-IE1 (164-584)-DsRed into BmN cells. Surprisingly, EGFP-IE (164-584) and IE1 (164-584)-DsRed co-localized in cells that had completed mitosis (Fig.6E). Taken together, these findings suggest that the N-terminus of BmIE1 might contain a sequence or a domain that is subject to degradation after mitosis.

### 2.5 NLSs identified here are conserved and can also function in a mammalian cell line

To determine whether NLSs we identified here can also function in a non-insect cell line, we constructed a plasmid in which the BmNPV *ie1* promoter was replaced with the CMV *ie1* immediate-early promoter so that fusion genes could be expressed in the NRK-52E rat kidney cell line. As can be seen from Fig.7, all NLSs were able to augment the nuclear targeting ability of EGFP-IE (Δ135-151), although to varying degrees. Moreover, the nuclear targeting abilities of the A3 fusion proteins were also improved by these NLSs (S10 Fig.). These results indicate that NLSs identified here are evolutionarily conserved.

**Fig. 7.**
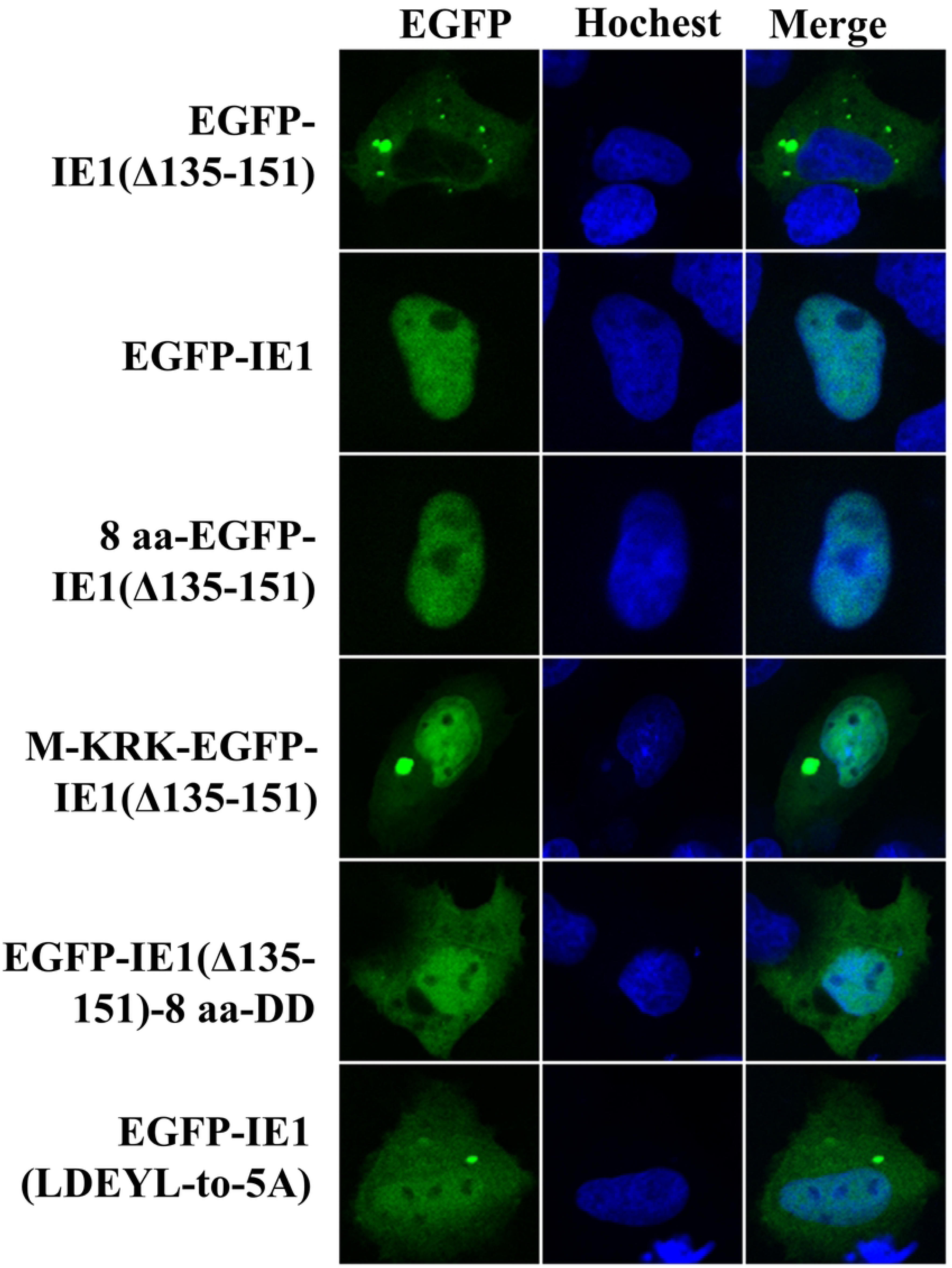
NLSs identified here are evolutionarily conserved. NRK-52E rat kidney cells were transfected with plasmids, fixed at 24 h p.t., stained with Hoechst 33342 and examined with a confocal microscope Magnification, X400.

### 2.6 A successful infection requires the nuclear levels of BmIE1 to reach certain thresholds

To determine whether the 29 aa bipartite NLS and the 8 aa monopartite NLS can be functionally complemented by others NLSs, we fused EGFP-IE1 (Δ135-151) and EGFP-IE1 with SV40 NLS and NLSs identified here. At an MOI of 0.05, vBmKO: SV40 NLS-EGFP-IE1 (Δ135-151) and vBmKO: M-KRK-EGFP-IE1 (Δ135-151) generated significantly lower titers than vBmKO: 8 aa-EGFP-IE1 (Δ135-151), vBmKO: EGFP-IE1, and vBmKO: 8 aa-EGFP-IE (Fig.8A). This result was consistent with the observation that, in plasmid-transfected BmN cells, SV40 NLS-EGFP-IE1 (Δ135-151) and M-KRK-EGFP-IE1 (Δ135-151) were also found in the cytoplasm, whereas the other fusion proteins localized completely to the nucleus (S11 Fig.). However, at an MOI of 5, they produced equivalent titers (Fig.8B). Notably, despite an additional copy of NLS at the N-terminus of 8 aa-EGFP-IE1, vBmKO: 8 aa-EGFP-IE1 did not differ statistically from vBmKO: EGFP-IE1 in terms of titers they produced at both MOIs tested (Fig.8A and 8B). Moreover, although EGFP-IE1 (Δ135-151) and EGFP-IE1 (Δ135-151)-8 aa both localized predominantly to the cytoplasm in plasmid-transfected BmN cells (Fig. 2D), unlike vBmKO: EGFP-IE1 (Δ135-151), vBmKO: EGFP-IE 1(Δ135-151)-8 aa was able to form plaques in its infected BmN cells. Addition of one or two Asp residues to the 8 aa domain significantly increased plaque diameters (Fig.8C). Taken together, these results indicate that differences in nuclear targeting abilities can be compensated by using high MOIs, and that if nuclear levels of BmIE1 fails to reach the lower threshold, no infection can be launched, whereas if the upper threshold is reached, the performance of BmIE1 can no longer be further improved by adding additional NLSs.

**Fig. 8.**
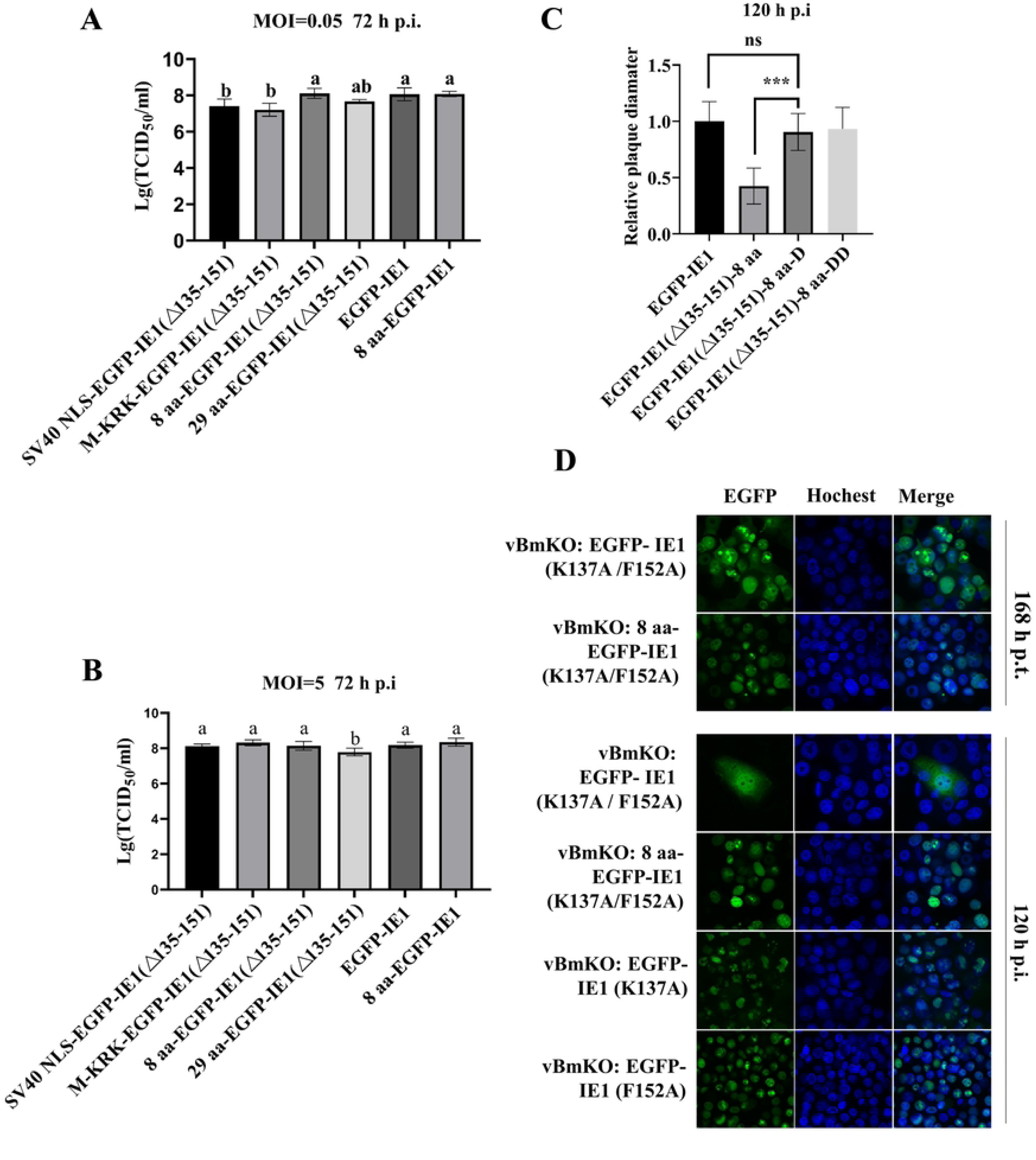
Effects of different NLSs on the titers of repaired vBmKO. Virus yields **A** (MOI=0.05) and **B** (MOI=5). Extracellular budded virus produced by BmN cells infected with vBmKO repaired by indicated by indicated fusion proteins was quantified at 72 h p.i. by TCID50 measurements. Reported values are averages ± standard deviations of virus yields from three independent infections. **C,** Effects of Asp attached to the C-terminus of the 8 aa monopartite NLS on the plaque diameters of vBmKO repaired by indicated fusion proteins. **D,** Infectivity of vBmKO repaired by EGFP-IE1 (K137A/F152A) and 8 aa-EGFP- IE1 (K137A/F152A) in bacmid-transfected and virus-infected BmN cells.

As the mutant IE1 (F152A) was functionally impaired, we asked whether a successful infection could be launched if we reduced its nuclear targeting ability by introducing the K137A mutation. As can be seen from Fig.8D, although vBmKO: EGFP- IE1 (K137A / F152A) was able to form plaques in bacmid-transfected BmN cells, no sign of infection was observed in its infected BmN cells. By contrast, vBmKO: EGFP- IE1 (K137A) and vBmKO: EGFP- IE1 (F152A) were both found to form plaques in their infected BmN cells. However, when we added an additional NLS (the 8 aa-EGFP NLS) to EGFP- IE1 (K137A / F152A), the plaque-forming ability was recovered for vBmKO: 8 aa-EGFP-IE1 (K137A / F152A). Consistently, using the mutant IE1 (Δ135-152), a similar result was observed. As can be seen from S12 Fig., in BmN cells transfected with bacmids: vBmKO: 8 aa-EGFP-IE1 (Δ135-152), vBmKO: 29 aa-EGFP-IE1 (Δ135-152), or vBmKO: SV40

NLS-EGFP-IE1 (Δ135-152), a successful infection was launched, as indicated by spreading fluorescent signals. However, in BmN cells infected with the supernatant from transfected BmN cells, a successful infection was observed only for vBmKO: 8 aa-EGFP-IE1 (Δ135-152) and vBmKO: 29 aa-EGFP-IE1 (Δ135-152), whereas no sign of infection was observed for vBmKO: SV40 NLS-EGFP-IE1 (Δ135-152). Taken together, these results further indicate that infection is not initiated until nuclear BmIE1 accumulates to a certain threshold.

### 2.7 The non-canonical NLS is sufficient for launching a successful infection

Why BmIE1 evolved 3 NLSs is intriguing. To answer this question, it is important to determine whether the non-canonical NLS (the Lys500 and Lys501 motif) alone is sufficient for launching a successful infection. In BmN cells transfected with the bacmid vBmKO: EGFP-IE1 (Δ135-137), infection was observed, as demonstrated by spreading fluorescent signals (S13 Fig.). Unexpectedly, in BmN cells infected with the supernatant from the transfected BmN cells, no fluorescent signal was observed. Interestingly, when EGFP was fused to the C-terminus of IE1 (Δ135-137), vBmKO: IE1 (Δ135-137)-EGFP was infectious both in transfected and infected BmN cells, although it generated significantly smaller plaques relative to vBmKO: IE1-EGFP (Fig.9A). Considering that the attached EGFP might have an effect on nuclear import, we also tested IE1 (Δ135-137) for its ability to rescue vBmKO. As can be seen from Fig.9B, the supernatant from BmN cells transfected with the bacmid vBmKO: IE1 (Δ135-137) was able to launch a successful infection, as indicated by rounding cells detached from the bottom of the plate.

**Fig. 9.**
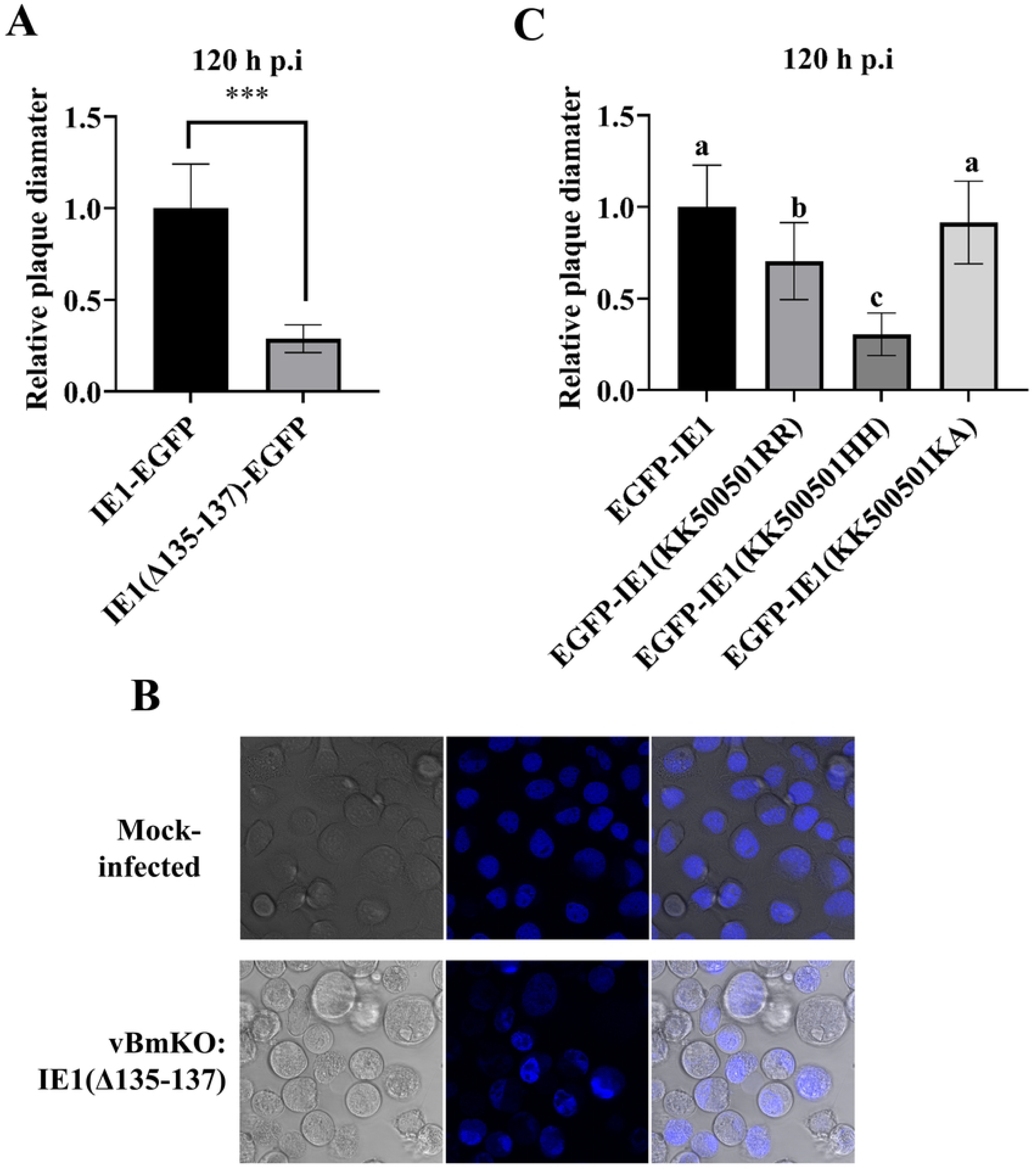
The non-canonical NLS alone is sufficient for initiating infection. **A,** The relative plaque diameters of vBmKO repaired by indicated fusion proteins. Twenty plaques were selected for each virus and their diameters were measured with Photoshop CS3. The ordinate shows the average diameter of vBmKO: IE1 (Δ135-137)-EGFP relative to that of vBmKO: IE1-EGFP. **B,** The infectivity of vBmKO repaired by IE 1(Δ135-137). **C,** Effects of mutations in the non-canonical NLS on the relative plaque diameters of vBmKO repaired by indicated fusion proteins. The ordinate shows the average diameter of each virus relative to that of vBmKO: EGFP-IE1.

We also evaluated the importance of Lys500 and Lys501 in the context of baculovirus infection. As the K500A mutation severely impaired the domain, we failed to observe plaques for the mutant. Therefore, the mutant was excluded in the plaque diameter analysis. As can be seen from Fig.9C, the K501A mutation appeared to have a minimal effect, and both the K500R/K501R mutant and the K500H/K501H mutant failed to fully complement the wild-type BmIE1. Plaque diameters appeared to be positively correlated with sequence non-specific binding abilities seen in mitotic cells (Fig. 5C). Taken together, these results indicate that Lys500 plays a major role in this domain and that it out-performs the other two basic residues.

### 2.8 Residues 57 to 151 are dispensable and the functional domains are separable and transferable

As BmIE1 is a multi-functional protein that contains multiple functional domains, we asked whether these domains were functionally separable and transferable. To this end, a series of IE1 mutants were constructed in which the domains are organized in various manners (Fig.10), and the infectivity of repaired vBmKO rescued by these mutants is summarized in Table 1.

**Fig. 10.**
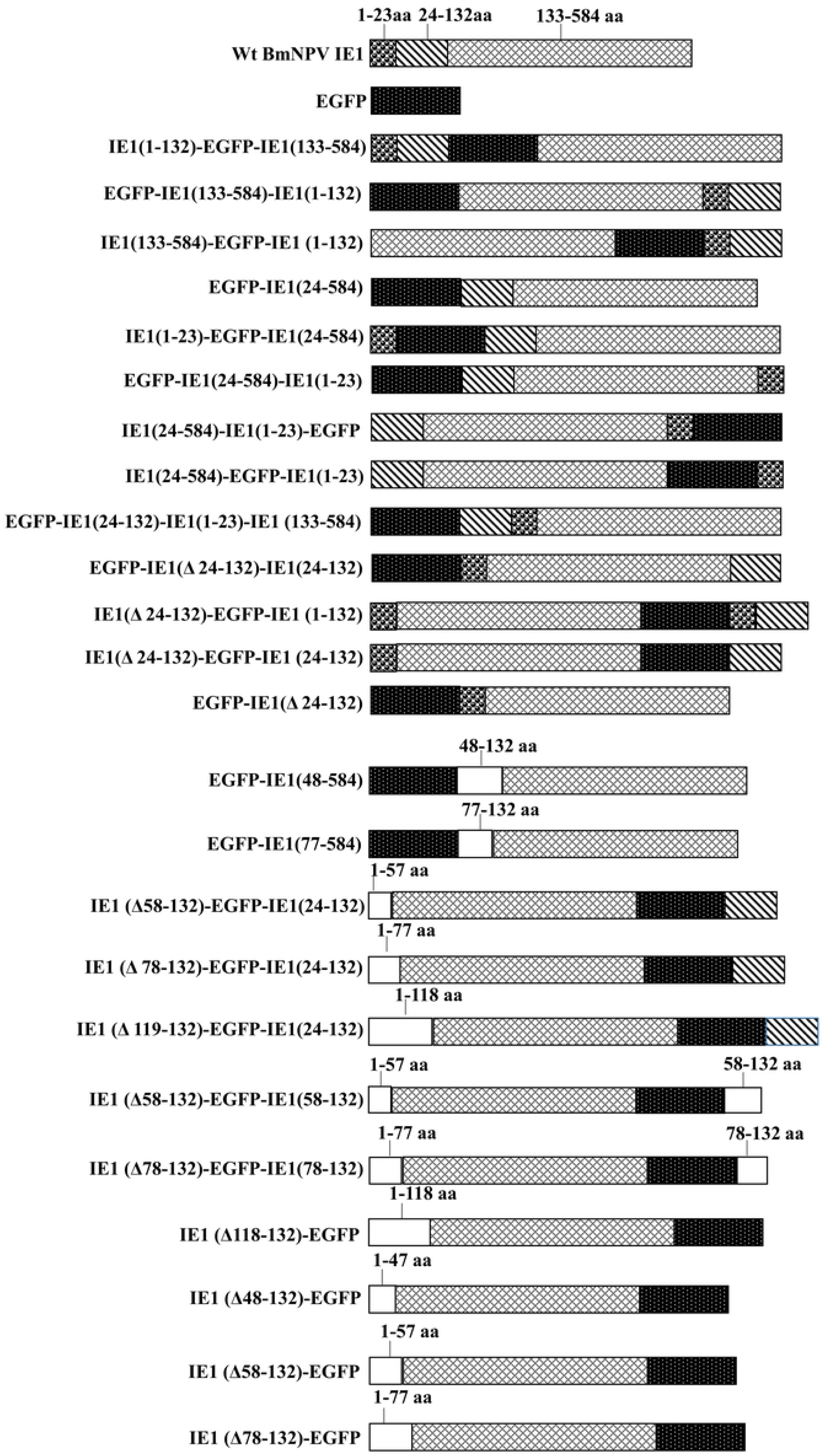
Schematic of reorganized BmIE1 and EGFP fusion proteins. BmIE1 are divided into 3 fragments, 1-23 aa, 24-132 aa and 133-584 aa, as indicated in the schematic. The white rectangles indicate fragments of BmIE1 and the residues they contain are shown above them.

**Table 1.**
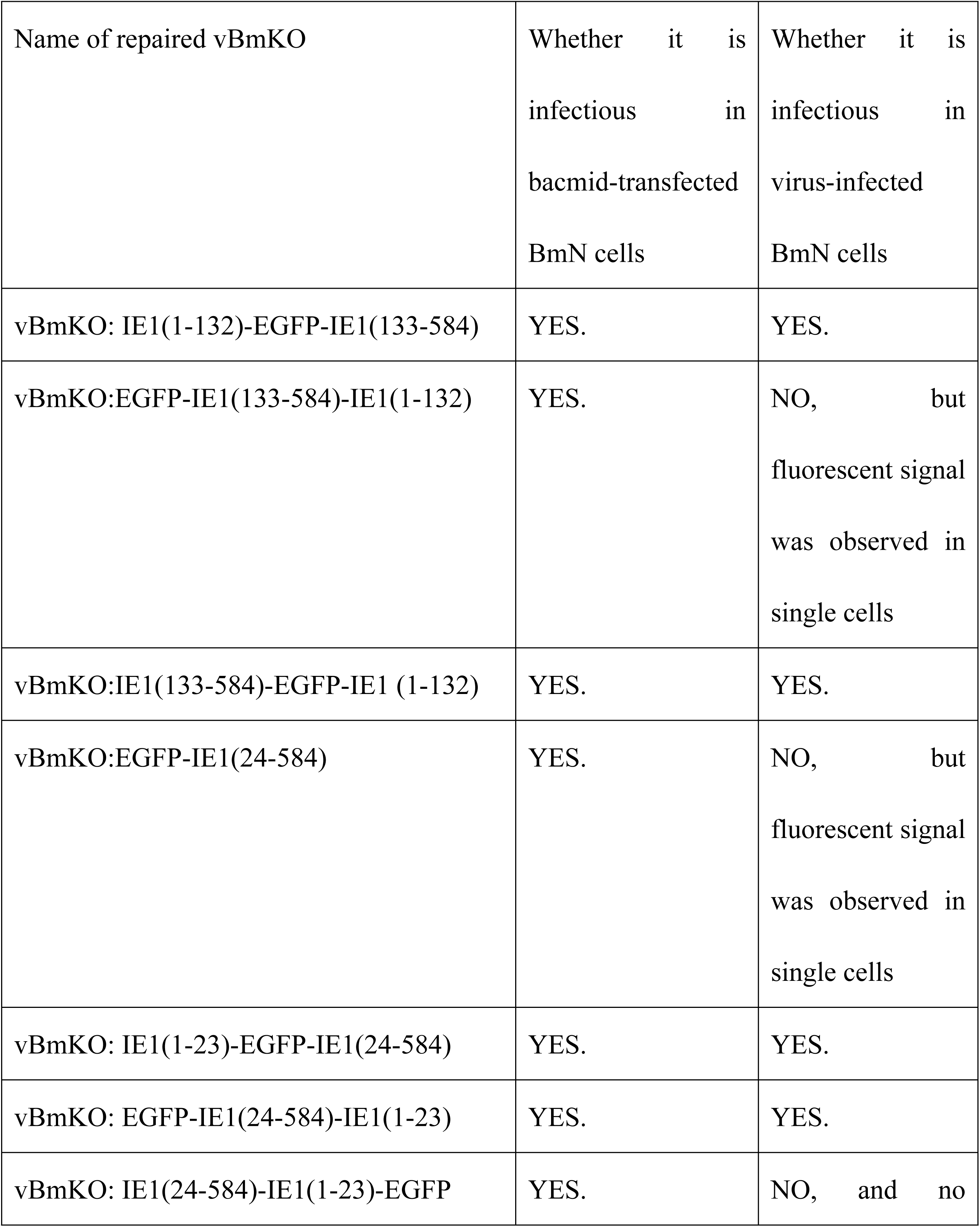
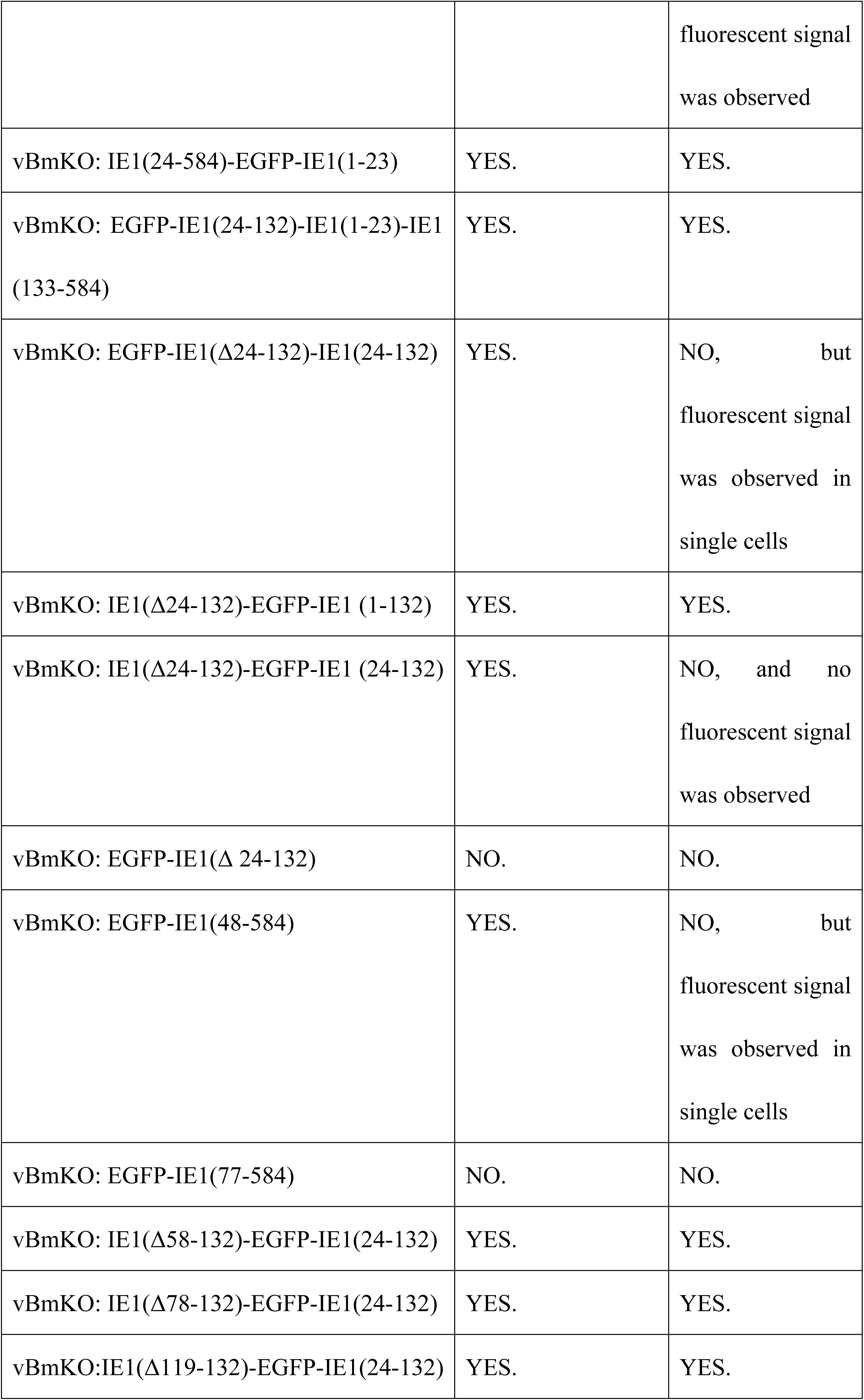
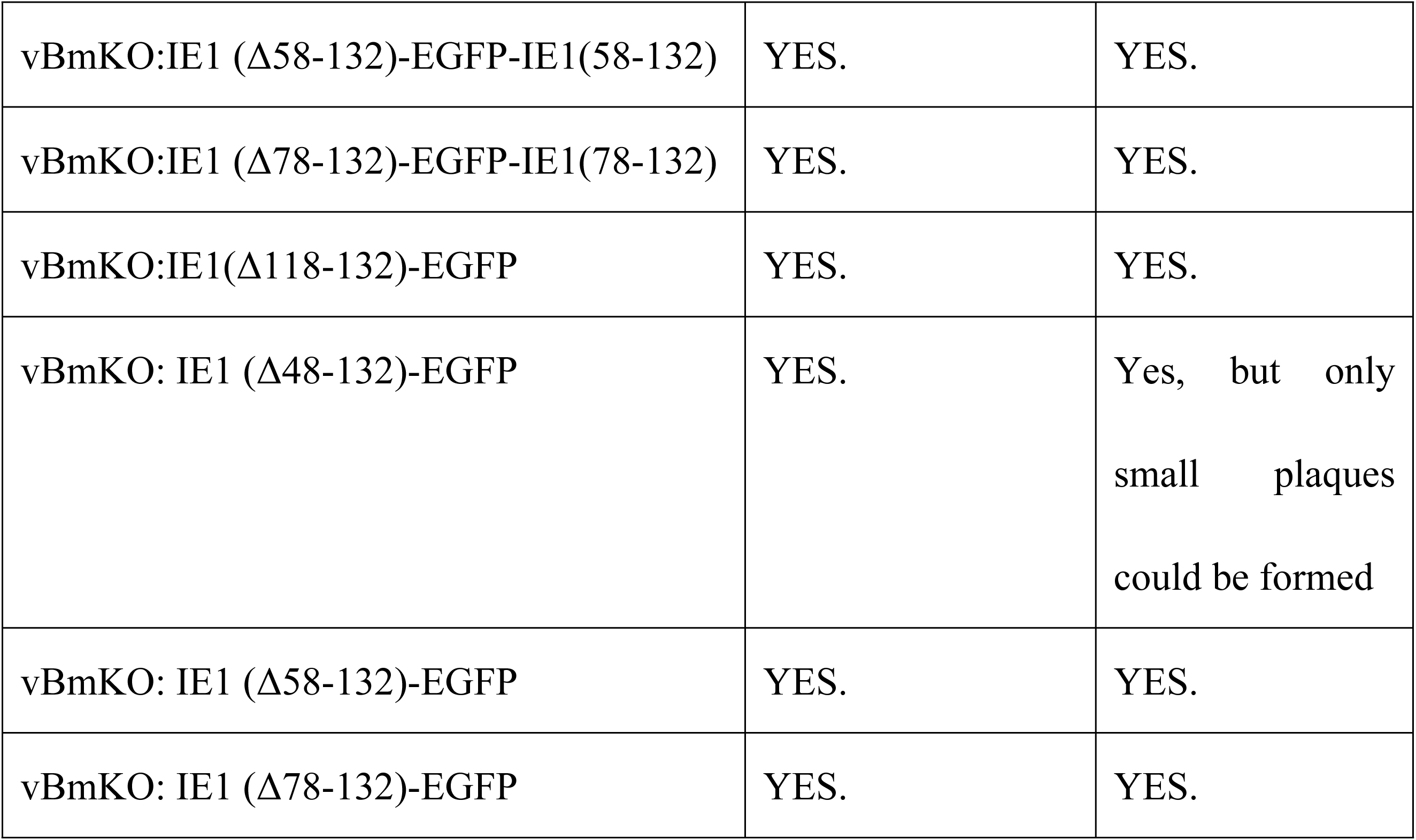
Infectivity of repaired vBmKO rescued by IE1 mutants in bacmid-transfected BmN cells and virus-infected BmN cells.

Based on the position of the KRK motif, we inserted EGFP at a position between Ala132 and Gly133 of BmIE1 to generate a fusion protein IE1 (1-132)-EGFP-IE1 (133-584). Surprisingly, insertion of EGFP at this position appeared to have a minimal effect on plaque diameters (Fig.11A). We next transferred IE1 (1-132) to the C-terminus of the fusion protein IE1 (133-584)-EGFP, and found that vBmKO: IE1 (133-584)-EGFP-IE1 (1-132) formed significantly smaller plaques than vBmKO: IE1 (1-132)-EGFP-IE1 (133-584) (Fig.11A), indicating that functional domains located in IE (1-132) appears to work better at the N-terminus of BmIE1.

The N-terminal 23 aa of AcIE1 has been shown to contain a replication domain and BmIE1 also possesses this domain at its N-terminus. Although vBmKO: EGFP-IE1 (24-584) was able to form plaques in bacmid-transfected BmN cells, it failed to form plaques in its infected BmN cells (Fig. 11B). To determine whether this replication domain is separable and transferrable, we inserted EGFP at a position between Asn23 and Gly24 of BmIE1, and we transferred IE1 (1-23) to the C-terminus of the fusion protein IE1 (24-584)-EGFP. Moreover, we transferred IE1 (1-23) to the C-terminus of the fusion protein EGFP-IE1 (24-584), and we also transferred IE1 (1-23) to a position between Ala132 and Gly133 of BmIE1. vBmKO rescued by these fusion proteins all formed plaques, and these plaques were not statistically different in diameter (Fig.11A). These results indicate that the 23 aa domain is indeed functional, separable and transferable. In support of this conclusion, vBmKO: IE1 (Δ24-132)-EGFP-IE1 (1-132), which has an extra copy of IE1 (1-23) at the N-terminus of the fusion protein, formed significantly larger plaques than vBmKO: IE1 (133-584)-EGFP-IE1 (1-132) (Fig.11A). However, vBmKO: IE1 (Δ24-132)-EGFP-IE1 (24-132) failed to form plaques in infected BmN cells (Fig.11B), indicating that the 23 aa domain at the N-terminus of the fusion protein alone is not sufficient for forming plaques. In support of this possibility, no sign of infection was observed in vBmKO: EGFP-IE1 (Δ 24-132)-transfected BmN cells (Table 1). Moreover, vBmKO: IE1 (1-132)-EGFP-IE1 (133-584) formed significantly larger plaques than the vBmKO: IE1 (1-23)-EGFP-IE1 (24-584) (Fig.11A). Taken together, these findings indicate that the 23 aa domain might be a functional part of the replication domain and that additional residues C-terminal to the 23 aa domain are required for the replication domain to exert a full activity.

**Fig. 11.**
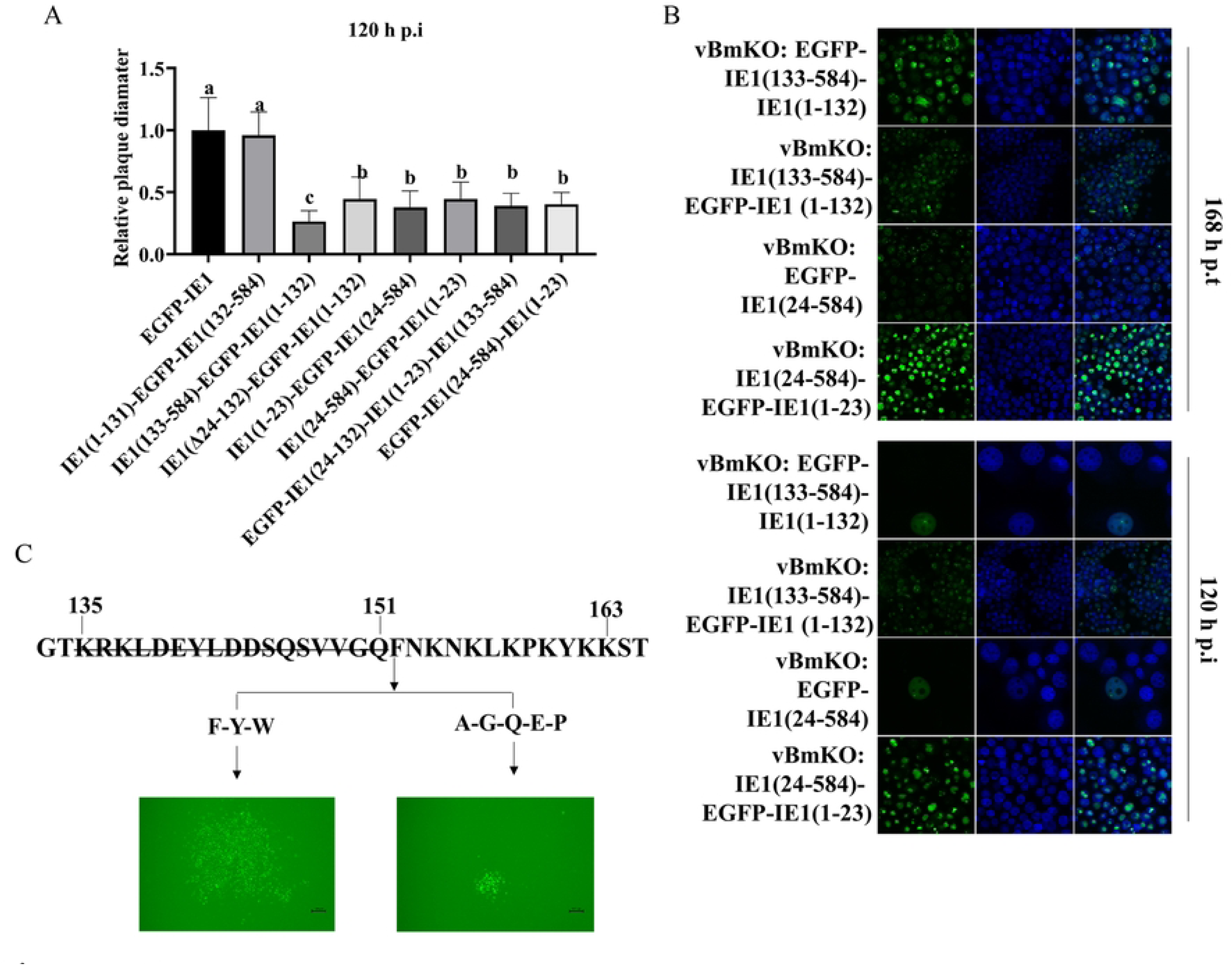
Effects of domain transfer and F152 substitution mutations on infectivity. **A,** Relative plaque diameter. Twenty plaques were selected for each virus repaired by indicated fusion proteins and their diameters were measured with Photoshop CS3. The ordinate shows the average diameter of each virus relative to that of vBmKO: EGFP-IE1. **B,** The transfection-infection assay. In bacmid-transfected BmN cells, signs of infection were observed at 168 h p.t., and in infected BmN cells, signs of infection were observed at 120 h p.i.. **C,** Effects of F152 substitution mutations on plaque diameters. In the point mutations, F152 was substituted for Y, W, A, G, Q, E, and P, and residues 135 to 151 were deleted, with the 8 aa-EGFP NLS at the N-terminus of the fusion proteins. Plaque diameter analysis was used to observe infectivity.

To test this possibility, we truncated 47 aa and 77 aa from the N-terminus of BmIE1, and found that, in bacmid-transfected BmN cells, as with vBmKO: EGFP-IE1 (24-584), vBmKO: EGFP-IE1 (48-584) was able to form plaques, whereas vBmKO: EGFP-IE1 (78-584) no longer formed plaques (Table 1). This finding indicate that, residues 24 to 77 can functionally complement residues 1 to 23.

To map those residues that can functionally complement residues 1 to 23, based on IE1 (Δ24-132)-EGFP-IE1 (24-132), we constructed internally-deleted mutants that retain more N-terminal residues including IE1(Δ58-132)-EGFP-IE1(24-132), IE1(Δ78-132)-EGFP-IE1 (24-132) and IE1 (Δ118-132)-EGFP-IE1 (24-132), and found that they were all able to rescue vBmKO in infected BmN cells. Next, we constructed IE1 (Δ58-132)-EGFP-IE1 (58-132), IE1 (Δ78-132)-EGFP-IE1 (78-132) and IE1 (Δ118-132)-EGFP, and vBmKO could be also recused by these mutants. Considering that IE1 (Δ118-132)-EGFP, in which residues 118 to 132 are deleted, was able to rescue vBmKO, we sought to identify those essential residues N-terminal to residue 118 of BmIE1. To this end, we constructed IE1 (Δ78-132)-EGFP, IE1 (Δ58-132)-EGFP, and IE1 (Δ48-132)-EGFP. Remarkably, all these mutants were able to rescue vBmKO in bacmid-transfected BmN cells, although vBmKO: IE1 (Δ48-132) could only form few small plaques in infected BmN cells. Taken together, these findings suggest that residues 58 to 132 of BmIE1 are dispensable for launching a successful infection in infected BmN cells.

Unlike vBmKO: SV40 NLS-EGFP-IE1 (Δ135-151), vBmKO: SV40 NLS-EGFP-IE1 (Δ135-152) failed to form plaques and no focal loci were observed in its infected BmN cells (S12 Fig.), indicating that residues 135 to 151 are dispensable and that Phe152 is a critical residue involved in hr binding. To confirm that Phe152 is vital for infection, we sought to determine whether Phe152 can be substituted for other residues. Given that sufficient IE1 nuclear import is also critical for a successful infection, we used the relatively strong 8 aa-EGFP NLS for the fusion proteins. Surprisingly, based on plaque diameter analyses, we observed that mutation of Phe152 to aromatic amino acids (i.e., Tyr and Trp) appeared to have little effect on plaque diameters, whereas mutation of Phe152 to other residues including Ala, Gly, Gln, Glu and Pro, seemed to have severe effects (Fig.11C). These results indicate that Phe152 might be involved in hr binding and that the benzene ring may play a pivotal role.

### 2.9 BmNPV could launch infection more efficiently in a BmN cell line over another BmN cell line by increasing transcription levels of immediate early genes

Interestingly, we observed that vBmKO: EGFP-IE1 could launch infection more efficently in a BmN cell line (small BmN cellss) over another BmN cell line (big BmN cells)(Fig. 12A). To dermine whether this is specific to vBmKO: EGFP-IE1, we also used a wild-type BmNPV bacmid virus that expressed EGFP under the control of the *ie1* promoter at the *polyhedrin* locus, and a similar result was observed (S14 Fig.). Notably, for vBmKO rescued by functionally impaired IE1, the difference was found to be even more significant. As can be seen from Fig.12B, fluorescent signal was observed at 24 h p.i. in more than half of the small BmN cells infected with 1 V of vBmKO: EGFP-IE1(K500H/K501H), whereas when the big BmN cells were infected with 1 V of the virus, fluorescent signal was observed in only a small number of cells even at 168 h p.i. and no sign of infection was observed, although when infected with 3 V or 10 V of the virus, signs of infection were observed at 168 h p.i.. These results indicate that the virus failed to infect the big BmN cells at a low MOI. To test this possibility, we used the two cell lines to determine the titers of vBmKO: EGFP-IE1 and vBmKO: SV40 NLS-EGFP-IE1(Δ135-151). Remarkably, for vBmKO: SV40 NLS-EGFP-IE1(Δ135-151), the titers determined by the big and small BmN cell lines were nearly 100-fold diffferent, and for vBmKO: EGFP-IE1, there was a 5.1-fold difference (Fig.12C). Consistently, vBmKO: EGFP-IE1 formed significantly larger plaques in the small BmN cells than in the big BmN cells (Fig.12D). Moreover, at an MOI of 0.05, the titer achieved at 5 d p.i. by vBmKO: EGFP-IE1 in the small BmN cells was nearly 2000-fold higher than the titer achieved by vBmKO: SV40 NLS-EGFP-IE1 (Δ135-151) in the big BmN cells (Fig.12E). Interestingly, vBmKO rescued by KRK-mutated IE1(with EGFP fused at its N-terminus) was able to launch infection in the bacmid-transfected small BmN cells (S13 Fig.), whereas no sign of infection or even no fluorescent signal was observed in the bacmid-transfected big BmN cells (Fig.4A). However, no fluorescent signal was observed in the small BmN cells infected with the supernatant from the bacmid-transfected small BmN cells.

**Fig. 12.**
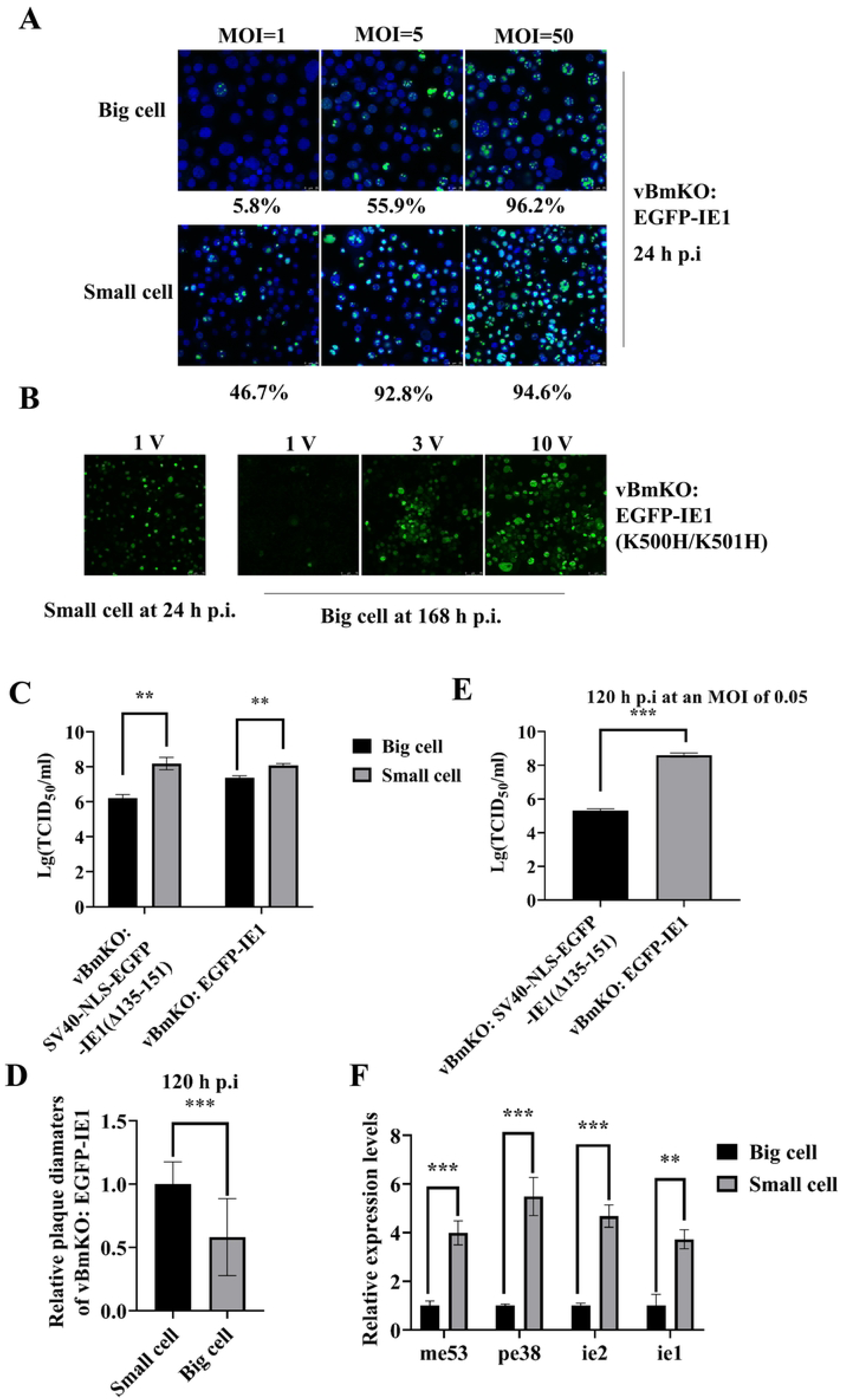
BmNPV could launch infection more efficiently in a BmN cell line over another BmN cell line by increasing transcription levels of immediate early genes. **A,** At 24 h p.i, fluorescent signal could be observed in higher proportions of the big BmN cells relative to the small BmN cells when the cells were infected with vBmKO:EGFP-IE1 at MOI=1 and MOI=5. **B,** vBmKO: EGFP-IE1 (K500H/K501H) failed to infect the big BmN cells at a relatively low MOI. **C,** Titer determination. The big BmN cells and the small BmN cells were used to determine the titers of vBmKO: SV40-NLS-EGFP-IE1(Δ135-151) and BmKO: EGFP-IE1 by TCID50 measurement. **D,** Relative plaque diameters of vBmKO: EGFP in the big and small BmN cells. Twenty plaques were selected for each group and their diameters were measured with Photoshop CS3. The ordinate shows the average diameter of vBmKO: EGFP-E1 in the big BmN cells relative to that of vBmKO: EGFP-IE1 in the small BmN cells. **E,** The big BmN cells were infected with either vBmKO: SV40-NLS-EGFP-IE1(Δ135-151) or BmKO: EGFP-IE1 at an MOI of 0.05 and extracellular budded virus produced was quantified at 120 h p.i. by TCID50 measurement. **F,** Relative expression levels of the immediate-early genes *me53*, *pe38*, *ie2* and *ie1* in the big BmN cells and the small BmN cells infected at an MOI of 5 with vBmKO: EGFP-IE1 at 2 h p.i. (** p <0.01 and *** p <0.001).

The two cell lines differ in size and the big BmN cells appear to attach more tightly to the bottom of the plate (S15 Fig.). To explore the molecular mechanism by which the small BmN cell line can more efficiently support BmNPV infection, we determined the expression levels of immediate-early genes at 2 h p.i. at an MOI of 5. As can be seen from Fig.12F, transcript levels of all immediate-early genes we tested were found to be significantly more abundant in the small BmN cells than in the big BmN cells. This finding indicate that enhanced expression of immediate-early genes might be critical for efficiently launching infection.

### 2.10 Multiple sequence alignment and phylogenetic analyses provide insight into when IE1 functional domains evolved

To gain insight into how ancient IE1 evolved, we employed multiple sequence alignment analysis of IE1 amino acid sequences by taking advantage of a rooted phylogenetic tree created using the concatenated amino acid sequence of all conserved genes among 64 publicly available baculovirus and nudivirus annotated genomes from species across alphabaculoviruses, betabaculoviruses, gammabaculoviruses, deltabaculoviruses, and nudiviruses[26]. As IE1 proteins are found only in alphabaculoviruses and betabaculoviruses, a total of 55 IE1 amino acid sequences were used in our alignment analysis. The alignment analysis revealed that IE1 amino acid sequences are more conserved in the C-terminus than in the N-terminus, with all the betabaculovirus IE1 amino acid sequences and IE1 amino acid sequences from two alphabaculoviruses (i.e., PeluSNPV and ApciNPV) lacking about 150 N-terminal residues found in the alphabaculoviruses. These results indicated that the additional N-terminal residues were acquired in the alphabaculovirus ancestor after it diverged from the betabaculovirus ancestor, and that the additional N-terminal residues were lost during evolution in some alpha II baculoviruses. Moreover, although basic domain I is found in both alpha I and alpha II baculoviruses, with no similar domain detected in the betabaculoviruses, it was less conserved in alpha II baculoviruses and was lost during evolution in some alpha II baculoviruses (Fig.13). Notably, the KRK motif is highly conserved in the alpha I baculoviruses but is found in only 3 species in the alpha II baculoviruses (Fig.13), indicating that the KRK motif was acquired in the alpha I baculovirus ancestor after it diverged from the alpha II baculovirus ancestor, and that the KRK motif found in the alpha II baculoviruses might be acquired through independent genetic events. The 8 aa-like monopartite NLS identified here was found only in four alpha I baculoviruses (i.e., AcMNPV, BmNPV, MmvNPV and TorNPV), and based on the phylogenetic tree[26], these viruses are most closely related, indicating that the 8 aa monopartite NLS was acquired by their common ancestor. Remarkably, the non-canonical NLS motif (K500 and K501) we mapped here was located in an extremely conserved domain in the C-terminus, with 3 residues (R495, W507 and H529) being identical across the 55 baculoviruses (S16 Fig.).

**Fig. 13.**
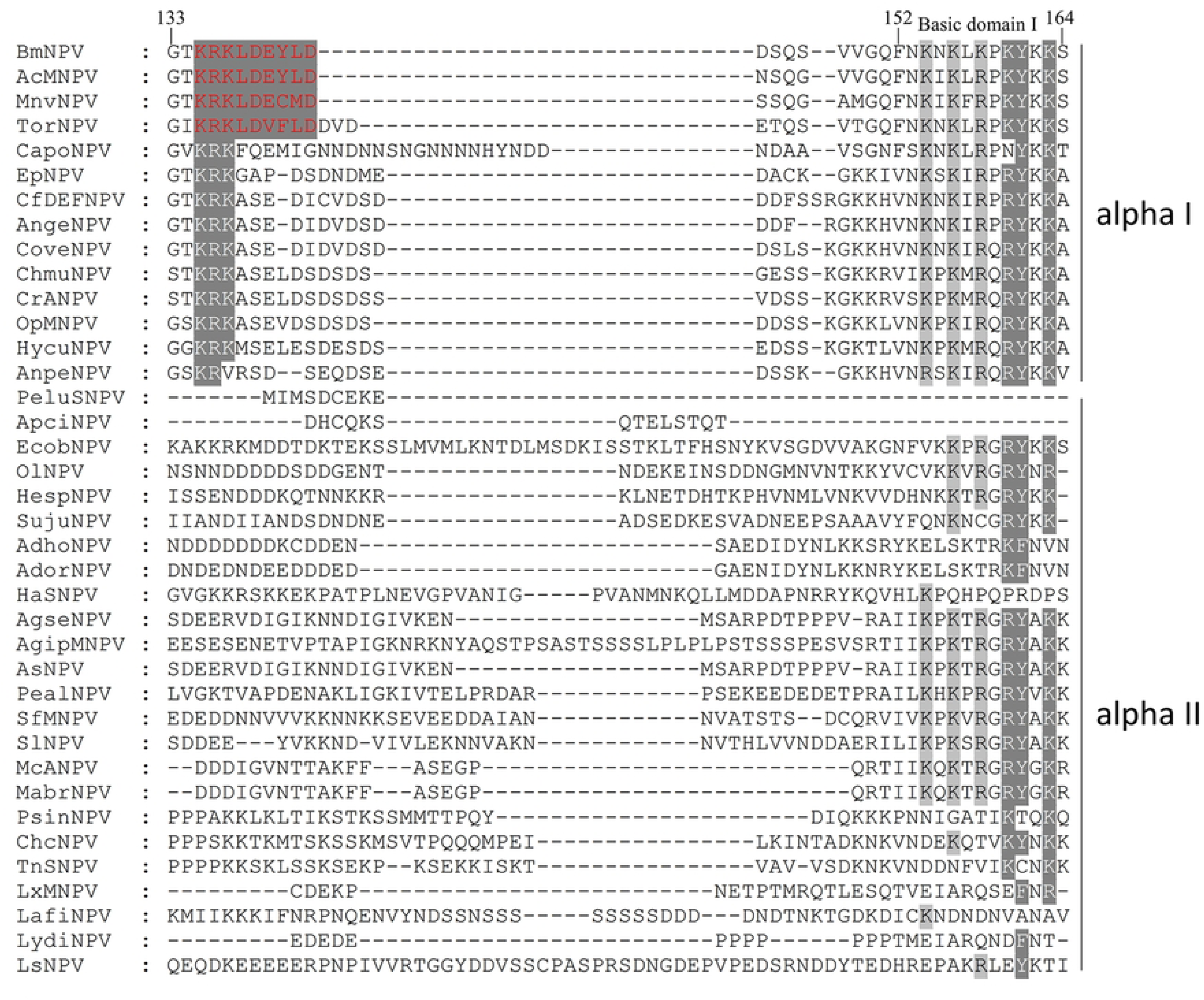
Analysis of amino acid sequence conservation of IE1 from alphabaculoviruses in the bipartite NLS. Conserved residues are shaded in grey. The 8 aa-like monopartite NLSs are marked in red.

## 3 Discussion

In the present study, we demonstrate that BmIE1 possesses a unique bipartite NLS that contains a monopartite NLS and thus explain why BmIE1 still localizes predominantly to the nucleus when the entire basic domain I is deleted. Moreover, we mapped a cryptic non-canonical NLS, which can function efficiently only when the size of the protein is proper, and we unraveled that this NLS is also involved in sequence non-specific DNA binding. Interestingly, we found that the 8 aa monopartite NLS can form a novel NLS when it is fused to the N-terminus of EGFP, and that even the motif M-KRK, when fused to the N-terminus of EGFP-IE1 (Δ135-151), can gain a nuclear targeting ability. We also revealed that N-terminally truncated BmIE1 can enter the nucleus through its sequence non-specific DNA binding ability. Notably, vBmKO: EGFP-IE1 (Δ135-137) failed to launch a successful infection in its infected BmN cells, whereas vBmKO: IE1 (Δ135-137)-EGFP was found to be able to spread in its infected BmN cells, indicating that the C-terminus of BmIE1 might form an NLS with EGFP. Inspection of the C-terminus of BmIE1 revealed a basic residue-rich sequence that is similar to basic domain I. Further investigation is needed to confirm this possibility. Our point mutation analyses showed that only Asp139 and Glu140 in the 8 aa monopartite NLS were allowed to be replaced with alanine and mutation of all the other residues resulted in a severe effect on nuclear targeting abilities (Fig.2C). By contrast, in the 29 aa bipartite, the KRK motif, which is shared by the 8 aa monopartite NLS, can be partially complemented by KRA, HHK or RRK (Fig.2A). Moreover, multiple sequence alignment analysis showed that basic domain I was the most conserved across baculoviruses, followed by the KRK motif and the 8 aa monopartite NLS (Fig.13). Based on these findings, we speculate that the bipartite NLS evolved after basic domain I and that the 8 aa monopartite NLS evolved after the bipartite NLS.

By using IE1 (164-584), an N-terminally truncated IE1 that lacks basic domain I, combined with the finding that EGFP-IE1 (164-584) can enter the nucleus through its binding to condensed chromosomes in mitotic cells (Fig.6B), we demonstrated that the sequence non-specific DNA binding ability of BmIE1 is independent of its hr binding ability. Moreover, EGFP-IE1(164-584)-(AVNAA) failed to bind to condensed chromosomes in mitotic cells (Fig.6B), and EGFP-IE1 (AVNAA) failed to form focal loci in bacmid-transfected BmN cells (Fig.1E), indicating that basic domain II is involved in the sequence non-specific DNA binding ability of BmIE1 on which the hr binding ability is dependent. Accordingly, we hypothesize that BmIE1 might bind to chromosomes through its sequence non-specific DNA binding ability and then slides on the chromosome until it encounters an hr sequence, which have been proposed to be capable of triggering a change in BmIE1 conformation that can result in accumulation of BmIE1 on the sequence[16]. In BmN cells transfected with pF-ie1p-EGFP-IE1 (AVNAA) and BmN cells transfected with vBmKO: EGFP-IE1 (AVNAA), a weak fluorescent signal was observed in the cytoplasm (Fig.1E), indicating that the nuclear targeting ability of basic domain II-mutated IE1 is slightly impaired. It is possible that mutation of KVNRR to AVNAA might result in a change in conformation that leads to a compromised nuclear targeting ability. Alternatively, the mutation might lead to the misfolding of the mutant and misfolded EGFP-IE1 (AVNAA) might result in formation of aggresomes in the cytoplasm [27, 28].

Although vBmKO: EGFP-IE1 (Δ135-137) were able to spread in bacmid-transfected BmN cell (S13 Fig.), no fluorescent signal was observed in its infected BmN cells. Moreover, due partially to an impaired nuclear targeting ability, in infected BmN cells, vBmKO: EGFP-IE1 (K137A/F152A) failed to spread, whereas vBmKO: 8 aa-EGFP-IE1 (K137A/F152A) regained infectivity (Fig.8D). As with vBmKO: EGFP-IE1 (Δ135-137), no fluorescent signal was observed in vBmKO: EGFP-IE1 (Δ135-151)-infected BmN cells, whereas vBmKO: EGFP-IE1 (Δ135-151)-8 aa was found to be able to spread in infected BmN cells despite the observation that EGFP-IE1 (Δ135-151)-8 aa still localized predominantly to the cytoplasm in plasmid-transfected BmN cells (Fig.8C and Fig.2D). All these findings suggest that sufficient levels of nuclear BmIE1 are critical for launching a successful infection. Furthermore, although vBmKO: EGFP-IE1 (24-584) were able to spread in bacmid-transfected BmN cells despite the deletion of the replication domain, only single focal locus was seen in infected BmN cells (Fig.11B), indicating that higher expression of functionally impaired BmIE1 can partially complement compromised functions. In support of this, due to lower expression of immediate-early genes in the big BmN cells than in the small BmN cells, when the big BmN cells were infected at a low MOI with vBmKO: EGFP-IE1 (K500H/K501H), fluorescent signal was observed in only a small number of cells even at 168 h p.i. and no sign of infection was observed (Fig.12B). By contrast, when the small BmN cells were infected with the same virus at the same MOI, fluorescent signal was observed at 24 h p.i. in more than half of cells (Fig.12B). Based on these findings, we hypothesize that, in the course of infection, the nuclear levels of BmIE1, or more precisely, the levels of BmIE1 total activity, should reach a total of 3 thresholds. Specifically, if threshold 1 (i.e., the lower threshold) is not reached, no fluorescent signal can be observed; if threshold 2 is not reached, fluorescent signal can only be observed in single cells; if threshold 3 (i.e., the upper threshold) is not reached, only small plaques can be formed; and if the the upper threshold is reached, an excess of BmIE1 total activity does not further increase infectivity. Consistent with this hypothesis, vBmKO: SV40 NLS-EGFP-IE1 (Δ135-152) failed to spread in infected BmN cells, with fluorescent signal observed only in single cells, whereas vBmKO: 8 aa-EGFP-IE1 (Δ135-152) and vBmKO: 29 aa -EGFP-IE1 (Δ135-152) were able to launch a successful infection in infected BmN cells (S12 Fig.). Moreover, lower infectivity of vBmKO: SV40 NLS-EGFP-IE1 (Δ135-151) and vBmKO: M-KRK-EGFP-IE1 (Δ135-151) could be compensated by a higher MOI, and the infectivity of vBmKO: 8aa-EGFP-IE1 was not increased despite an additional 8 aa-EGFP NLS at the N terminus relative to vBmKO: EGFP-IE1 (Fig.8A and 8B). Interestingly, a study investigating AcMNPV DNA polymerase (Ac DNApol) reported that an Ac DNApol–null virus, which was repaired with a mutated Ac DNApol, was able to spread in bacmid-transfected Sf-21 cells, but failed to spread in infected Sf-21 cells[29]. However, it was thought that virus production may have occurred but were below the level of detection[29]. Based on our findings, we speculate that the repaired virus might behave like vBmKO: EGFP-IE1 (Δ135-137), in which EGFP-IE1 (Δ135-137) is incapable of reaching threshold 1 in infected BmN cells. Moreover, our finding with the big BmN cells and the small BmN cells may offer an explanation for studies investigating AcMNPV orf34 (Ac34). Ac34 was shown to be essential in one study[30], however, another study using the same deletion strategy showed that infectious BVs could be still detected and found that, when the transfection efficiency was low, infectious BVs was hard to be detected using 50% tissue culture infective dose endpoint dilution assay[31]. In the present study, we found that, when the big BmN cells were transfected with vBmKO: EGFP-IE1 (Δ135-137), no fluorescent signal was observed or fluorescent signal was observed only in single cells (Fig.4A), depending on the transfection efficiency, whereas when the small BmN cells were transfected with the same bacmid, fluorescent signal was easily observed or signs of infection were observed if the transfection efficiency was high (S13 Fig.). Accordingly, we speculate that Ac34 might play a non-essential role which can be partially complemented by higher expression of viral genes and that different results of the studies might result from the cell lines they used.

Based on the finding that both vBmKO: IE1 (133-584)-EGFP-IE1 (1-132) and vBmKO: IE1 (1-132)-EGFP-IE1 (133-584) could formed plaques, with the latter forming significantly larger plaques than the former (Fig. 11A), we hypothesize that IE1 might have evolved the 132 aa residues through horizontal gene transfer and gene fusions, and that ancient IE1 without the 132 aa residues might cooperate with proteins that possessed functional domains that are contained in the 132 aa residues through protein-protein interactions. Given that IE1 contains multiple domains in the 132 aa residues, it is possible that these genetic events might have occurred multiple times. Consistent with this hypothesis, the 23 aa replication domain was also shown to be transferrable (Fig.11A and B), although it failed to exert a full activity due to the exclusion of the additional 34 residues (residues 24 to 57), which were shown in the present study to be critical for launching a successful infection (Table 1). Multiple sequence alignment analyses also revealed that IE1 amino acid sequences from betabaculoviruses appear to lack the 132 aa residues. Moreover, the hypothesis is further supported by the findings that residues 135 to 151 are shown to be dispensable and that vBmKO: IE1 (Δ135-151), in which the 29 aa bipartite NLS and the 8 aa monopartite NLS are deleted, was still able to launch a successful infection (Fig.9A and B).

The threshold hypothesis we propose here for IE1 may explain why ancient IE1 had evolved two additional NLSs, screened out the best residues and acquired multiple domains. Specifically, although vBmKO: IE1 (Δ135-151)-EGFP was able to launch a successful infection, it could only form significantly smaller plaques than vBmKO: IE1–EGFP due to a much weaker nuclear targeting ability (Fig.9A). IE1 with a stronger nuclear targeting ability can reach threshold 3 much faster and thus initiates an infection faster. Moreover, the finding that vBmKO: EGFP-IE1 (K137A/F152A) failed to spread in infected BmN cells, whereas vBmKO: 8 aa-EGFP-IE1 (K137A/F152A) regained infectivity (Fig.8D), also suggests that a stronger nuclear targeting ability can compensate for deleterious mutations in functional domains. This is important for IE1 molecular evolution because a stronger nuclear targeting ability means more leeway for IE1 to screen out the best residues. Indeed, our substitution analyses showed that IE1 had screened out the best residues for the KRK motif, residue 152 and the non-canonical NLS. Fusing domains that are functionally related and organizing them in a proper manner normally can achieve a higher total activity than that achieved by multiple proteins connected through protein-proteins interactions. The finding that vBmKO: IE1 (1-132)-EGFP-IE1 (133-584) could form significantly larger plaques than vBmKO: IE1 (133-584)-EGFP-IE1 (1-132) demonstrates that, even being fused together, the same domains when organized in a different manner can lead to differing activities (Fig.11A).

The finding that, due to lower expression of its immediate-early genes in the big BmN cells than in the small BmN cells, vBmKO: EGFP-IE1 (K500H/K501H) failed to infect the big BmN cells at a low MOI (Fig.12B), offers another explanation for IE1 molecular evolution. Host cells might have driven the molecular evolution of IE1 by screening out viruses that possessed an IE1 that had an improved nuclear targeting ability, that had selected the best residues for important domains and that had acquired related functional domains. Viruses with an improved IE1 could have a better efficiency of infecting host cells, could have the potential to infect more cell types or host species, and thus could have more leeway for genetic events. Therefore, it is the unique feature of threshold-based accumulation of IE1 total activity that drove the evolution of ancient IE1. The molecular evolution of IE1 was a process of IE1 screening out the best residues involved in functional domains (e.g., nuclear import and DNA binding), acquiring multiple NLSs and acquiring multiple domains through gene transfer and gene fusion to reach infection-initiating thresholds at a faster pace.

## 4 Materials and methods

### 4.1 Plasmids, bacmids and cells

Plasmids used in this study were constructed using either the restriction enzyme strategy or the seamless cloning strategy and all constructs were verified by sequencing, and primers utilized are listed in S1 Table. The BmNPV bacmid was purchased from Shanghai LMAI Bio Co., ltd. An *ie1*-null bacmid (vBmKO) was generated using a selection-marker free method, which was previously described [12]. Two *Bombyx mori* cell lines BmN (originated from ovary), i.e., the big BmN cells and the small BmN cells, were preserved in our laboratory and maintained at 27 °C in TC-100 medium (Sigma) supplemented with 10% fetal bovine serum (FBS).

### 4.2 Transfection

BmN cells were seeded onto a 12-well plate 12 h prior to transfection. Monolayers were overlaid with transfection mix containing cellfectin (Gibco) and plasmid or bacmid DNA in TC-100 medium and were then incubated at 27 °C for 4-6 h. After the incubation, the transfection mix was replaced with supplemented TC-100 medium and cells were then cultured at 27 °C.

### 4.3 Plaque assay

A monolayer of BmN cells in a 6-well plate were infected with viruses at a low MOI and the inoculum was replaced with 1% low-melting agarose dissolved in supplemented TC-100 at 2 h p.i.. At 120 h p.i., viral plaques were photographed using a fluorescence microscope and their diameters were measured using Adobe Photoshop CS3.

### 4.4 Quantitative real-time PCR

The big and small BmN cells were infected with vBmKO: EGFP-IE1 at an MOI of 5. At 2 h p.i., total RNA was extracted using a Qiagen kit, digested by RNase-free DNase, and then reversed transcribed into cDNA. RT-PCR was performed on optical bio-Raid 96 using SYBR Green for immediate-early genes including *ie1*, *ie2*, *me53* and *pe38*. The primers for the house-keeping gene, silkworm translation initiation factor 4A (sw22934), which was used to normalize gene expression, and the primers for the immediate-early genes, were described elsewhere [18].

### 4.5 Confocal microscopy

The transfected cells were washed three times with 1 ml of phosphate-buffered saline (PBS) at 48 h p.t., fixed by 4% paraformaldehyde, and then stained with Hoechst 33342. Cells were imaged on a Leica SP5 confocal laser scanning microscope (CLSM) with a 40X lens.

### 4.6 Statistical analysis

The statistical analyses were performed using IBM SPSS Statistics 26. Student’s t-test was employed to determine statistical significance. Values are presented as the mean ± SD from at least three independent biological replicates.

## Acknowledgements

We are grateful to Prof. Shiqing Xu and Prof. Xiaolong Hu for providing the BmN cell lines. We thank Xiaoxia Zhang and Mei Yin for their assistance in confocal microscopy.

## Author contributions

**Wujie Su:** Conceptualization, Methodology, Experimental design and implementation, Writing - original draft. **Chaoli Fang:** Resources, Writing-review. **Jiru He:** Resources, Writing - review. **Wenbing Wang:** Resources, Writing- review & editing. **Fanchi Li:** Resources, Writing - review & editing. B**ing Li:** Supervision, Project administration, Writing - review & editing.

## Data Availability

All data included in this study are available upon request by contacting the corresponding author.

## Ethics approval

This study did not require ethical approval.

## Consent to participate

Not applicable.

## Consent to publish

Not applicable.

## Supporting information captions

**S1 Fig. Analysis of amino acid sequence conservation of IE1 between BmIE1 and AcIE1.** Different residues between BmIE1 and AcIE1 are shaded in black.

(TIF)

**S2 Fig. Subcellular localization of N-terminally and C-terminally truncated IE1 in plasmid-transfected BmN cells.** BmN cells were transfected with plasmids, fixed at 48 h p.t., stained with Hoechst 33342 and examined with a confocal microscope Magnification, X400.

(TIF)

**S3 Fig. Subcellular localization of P143 fused with NLSs identified in the present study.** BmN cells were transfected with plasmids, fixed at 48 h p.t., stained with Hoechst 33342 and examined with a confocal microscope Magnification, X400.

(TIF)

**S4 Fig. Subcellular localization of BmIE1 with point mutations in the residues 135 to 163.** BmN cells were transfected with plasmids, fixed at 48 h p.t., stained with hoechst 33342 and examined with a confocal microscope Magnification, X400.

(TIF)

**S5 Fig. Subcellular localization of IE1 mutants in which basic domain I and residues N-terminal to it are deleted.** BmN cells were transfected with plasmids, fixed at 48 h p.t., stained with Hoechst 33342 and examined with a confocal microscope Magnification, X400.

(TIF)

**S6 Fig. Subcellular localization of IE1 mutants in which the 8 aa monopartite NLS is attached with different lengths of EGFP N-terminal residues.** BmN cells were transfected with plasmids, fixed at 48 h p.t., stained with Hoechst 33342 and examined with a confocal microscope Magnification, X400.

(TIF)

**S7 Fig. Subcellular localization of IE1 mutants in which point mutations are introduced to the 8 aa-EGFP NLS in the 8 aa domain as well as variants of M-KRK-EGFP.** BmN cells were transfected with plasmids, fixed at 48 h p.t., stained with Hoechst 33342 and examined with a confocal microscope Magnification, X400.

(TIF)

**S8 Fig. Subcellular localization of IE1 mutants in which the 8 aa monopartite NLS is deactivated and the linker lengths of the bipartite NLS are varied.** BmN cells were transfected with plasmids, fixed at 48 h p.t., stained with Hoechst 33342 and examined with a confocal microscope Magnification, X400.

(TIF)

**S9 Fig. Loss of sequence non-specific DNA binding abilities resulting from K422V/K423D, KKVKK-to-AAVDA and K500V/K5001D mutations in mitotic cells.** BmN cells were transfected with plasmids, fixed at 48 h p.t., stained with Hoechst 33342 and examined with a confocal microscope Magnification, X400.

(TIF)

**S10 Fig. NLSs identified here are evolutionarily conserved and can increase the nuclear targeting ability of the fusion protein EGFP-A3.** NRK-52E rat kidney ells were transfected with plasmids, fixed at 24 h p.t., stained with Hoechst 33342 and examined with a confocal microscope Magnification, X400.

(TIF)

**S11 Fig. Subcellular localization of EGFP-IE1 (Δ135-151) and EGFP-IE1 that are fused with NLSs indentified here.** BmN cells were transfected with plasmids, fixed at 48 h p.t, stained with Hoechst 33342 and examined with a confocal microscope Magnification, X400.

(TIF)

**S12 Infectivity of vBmKO repaired by 8 aa-EGFP-IE1(Δ135-152), 29 aa-EGFP-IE1(Δ135-152) and SV40 NLS-EGFP-IE1(Δ135-152) in bacmid-transfected and virus-infected BmN cells.**

(TIF)

**S13 Fig. Infectivity of vBmKO repaired by IE1 mutants in which the bipartite NLS and the monopartite NLS are impaired in the bacmid-transfected small BmN cells.**

(TIF)

**S14 Fig. Infectivity of wild-type EGFP-expressing BmNPV bacmid viruses in the big BmN cells and the small BmN cells.** Both cell lines were infected with the virus at MOIs of 1, 5 and 50, fixed at 24 h p.i, stained with Hoechst 33342 and examined with a confocal microscope Magnification, X400.

(TIF)

**S15 Fig. Images of the big BmN cells and the small BmN cells.** The cells were examined with a confocal microscope. Magnification, X400.

(TIF)

**S16 Fig. Analysis of amino acid sequence conservation of the C-terminus of IE1 proteins among 56 baculoviruses.** Extremely conserved residues are shaded in black and moderately conserved ones in grey.

(TIF)

**S1 Table. Primers used in the study**

